# Genome-wide *in vivo* dynamics of cohesin-mediated loop extrusion and its role in transcription activation

**DOI:** 10.1101/2024.10.02.616323

**Authors:** Tessa M. Popay, Ami Pant, Femke Munting, Morgan E. Black, Nicholas Haghani, Jesse R. Dixon

## Abstract

The organization of the genome in three-dimensional space is highly dynamic, yet how these dynamics are regulated and the role they play in genome function is poorly understood. Here, we utilized acute depletion of NIPBL to characterize the role of cohesin-mediated loop extrusion *in vivo*. Using this approach, we found that many chromatin loops are rapidly diminished upon loss of NIBPL, consistent with recent single locus imaging studies showing that chromatin loops are transient. However, we also identified cohesin-dependent chromatin loops that are associated with distinct chromatin states and may be “long-lived”, given that they require NIPBL for their establishment upon mitotic exit, but are persistent when NIPBL is depleted from interphase cells. In addition to the reformation of 3D genome structures, mitotic exit coincides with widespread transcriptional activation. We found that NIPBL is essential for establishing the expression of lineage-defining genes during the M-G1 transition but has diminished impact on the steady-state maintenance of their expression. At genes sensitive to its depletion, NIPBL supports a unique local genome organization defined by greater spatial proximity to nearby super-enhancers and weaker transcription start site insulation of genomic contacts. Overall, we show that NIPBL-mediated loop extrusion is critical to genome organization and transcription regulation *in vivo*.

## Introduction

In mammals, the genome is organized into hierarchical structures across three-dimensional space. This organization has been reported to play important roles in transcription regulation and DNA repair, with its dysfunction particularly associated with developmental disorders and cancer (*1, 2*). At the sub-chromosomal level, 3D genome organization is comprised of chromatin loops and topologically associating domains (TADs), which are thought to form through the process of loop extrusion driven by the interplay of cohesin and CTCF (*3*–*6*). Current models for cohesin-mediated loop extrusion suggest this process can be highly dynamic, yet we understand relatively little about *in vivo*, genome-wide cohesin dynamics and the influence they have on the regulation of transcription.

Early efforts to study cohesin dynamics *in vivo* using global ATP depletion led to the conclusion that chromatin loops are stable (*7*), yet recent live-cell imaging approaches at single loci have suggested that chromatin loops are able to persist for only 10-20 minutes (*8, 9*). The transient nature of chromatin loops is consistent with the ∼20 minute residence time of most cohesin on chromatin during G1 (*10*). While the longevity of chromatin loops has not been characterized genome-wide, the transient association of cohesin with chromatin suggests that the majority of chromatin contacts it facilitates are similarly transient. *In vitro*, cohesin-mediated loop extrusion requires the accessory protein NIPBL, which was originally described as a cohesin loading factor (*11, 12*). NIPBL likely facilitates loop extrusion through its ability to activate the ATPase activity of cohesin (*13*). However, the contribution of NIPBL to the binding of cohesin to chromatin and to 3D genome organization is under-explored in mammalian cells (*14*–*16*), particularly at the shorter time-scales that are relevant to understanding chromatin looping dynamics.

To address these questions, we established a system for the acute depletion of NIPBL to disrupt chromatin looping dynamics across multiple phases of the cell cycle. While many loops were rapidly lost upon NIPBL depletion, we found that a subset of cohesin-dependent loops persisted in the absence of NIPBL. Notably, these loops require NIPBL for their establishment during mitotic exit, suggesting distinct roles of NIPBL in the formation versus maintenance of different populations of chromatin loops *in vivo* and highlighting the possibility that chromatin loop longevity may vary genome-wide. Furthermore, given the essential role of the core cohesin complex in sister chromatid cohesion (*17*) and a dispensability for NIPBL for cohesion during mitotic exit, we used NIPBL depletion to test the role of post-mitotic genome reorganization in transcription reactivation (*18, 19*). We found that cohesin-mediated loop extrusion is particularly critical for establishing the expression of super-enhancer linked genes during mitotic exit. Taken together, our data illuminates the contribution of NIPBL to cohesin-mediated loop extrusion dynamics *in vivo*, and reveals how disruption to these dynamics affects the regulation of transcription.

## Results

### Acute depletion of NIPBL from hTERT RPE-1 cells

To better understand genome-wide dynamics of cohesin-mediated loop extrusion *in vivo*, we employed the dTAG system for acute depletion of NIPBL (*20, 21*). Using CRISPR/Cas9 homology-directed repair, we knocked-in an FKBP12^F36V^ tag to the N-terminus of NIPBL in the immortalized hTERT RPE-1 cells, generating two independent clonal cell lines expressing either P2A-linked mCherry (NIPBL-A2) or mGreenLantern (NIPBL-D7) (**Figure S1A-C**). Degradation of HA-NIPBL in both of these clones was rapid, beginning within one hour of treatment with dTAG^V^-1 (**Figure 1A, Figure S1D**). Previous studies have shown that NIPBL disruption leads to degradation of its heterodimer partner MAU2 and this has been used as an independent measure of NIPBL loss (*22, 23*). Upon treatment with dTAG^V^-1, MAU2 was depleted within 3 hours (**Figure 1A, Figure S1D**), confirming that our degradation of NIPBL is complete. The core cohesin subunits SMC3 and RAD21 remained largely stable with dTAG^V^-1 treatment (**Figure 1A, Figure S1D**). These observations support the use of the NIPBL-A2 and NIPBL-D7 cell lines for investigation into the role of NIPBL in regulating loop extrusion *in vivo*. We additionally engineered control cells for acute depletion of RAD21 using the dTAG system in the same hTERT RPE-1 background (**Figure S1E-G**) to compare the consequences of perturbing all cohesin-dependent genome organization with disrupting the formation of chromatin loops via extrusion.

**Fig. 1:**
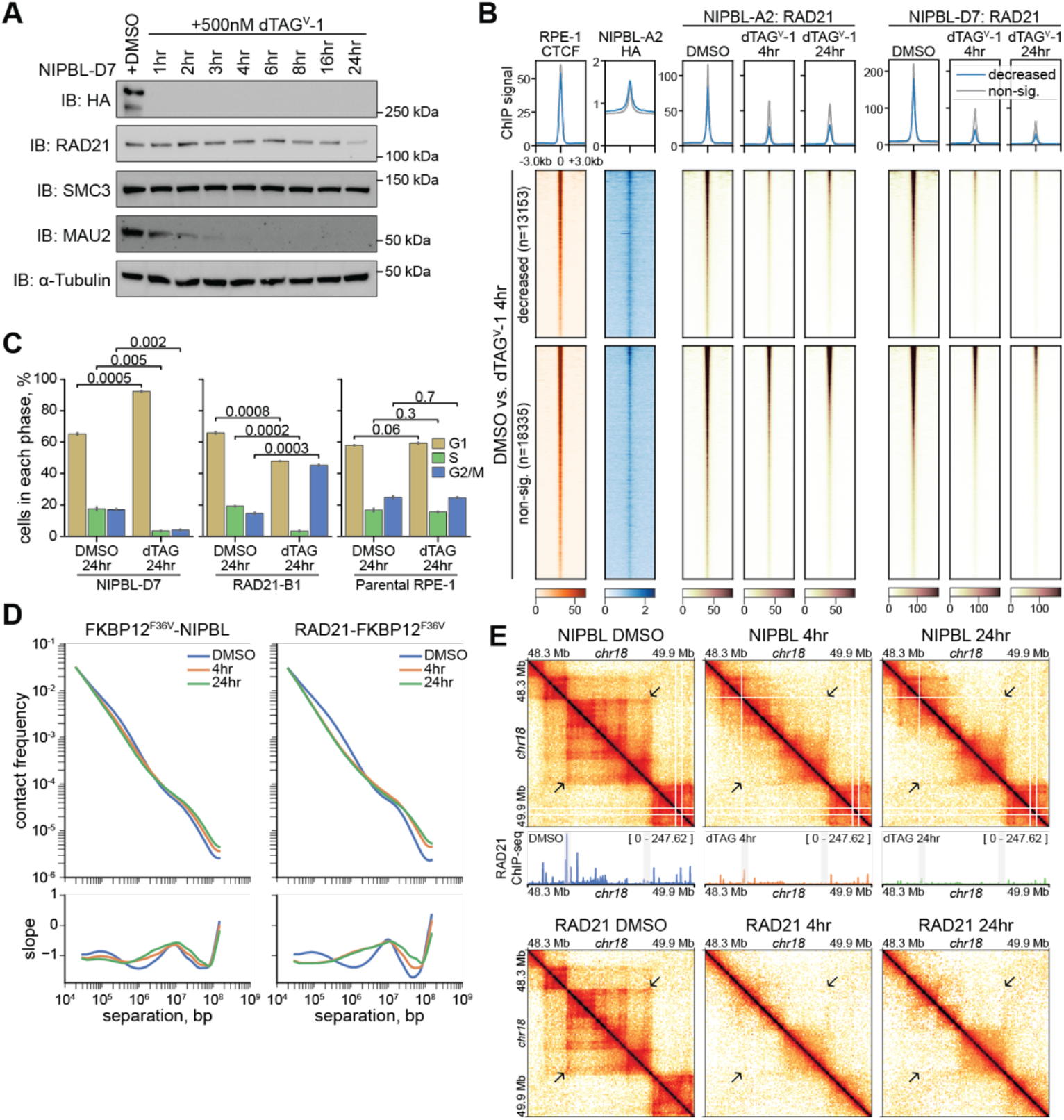
Acute depletion of NIPBL results in G1 cell cycle arrest and disruption to 3D genome organization. (**A**) NIPBL-D7 cells were treated with DMSO for 24 hours or dTAG^V^-1 for varying lengths of time, with the rate of HA-NIPBL depletion determined by western blotting. Also shown are the effects of NIPBL depletion on core cohesin subunits RAD21 and SMC3, and the cohesin accessory protein MAU2. α-Tubulin is shown as a loading control. (**B**) Scale factor-normalized RAD21 ChIP signal at significantly decreased and non-significantly changed peaks from differential analysis of RAD21 ChIP-seq with a 4-hour dTAG^V^-1 treatment in FKBP12^F36V^-NIPBL cells. Additionally shown is RPKM-normalized ChIP signal for HA-NIPBL and CTCF. RAD21 ChIP-seq from NIPBL-A2 and NIPBL-D7 cell lines were treated as replicates for differential analysis. (**C**) Quantification of cell cycle distribution by propidium iodide staining of NIPBL-D7 (left), RAD21-B1 (middle), and parental RPE-1 cells treated with DMSO or dTAG^V^-1 for 24 hours, as measured by flow cytometry. Independent, two-sided t-tests were performed on cell cycle distributions for each cell line using two replicates. Example distributions are shown in Figure S2F. (**D**) Relationship between distance and contact frequency at a 10 kb resolution based on Hi-C performed in cells depleted of NIPBL (left) or RAD21 (right) for 4 or 24 hours. To account for the accumulation of 4N cells with RAD21 depletion, DMSO- and dTAG^V^-1-treated nuclei were first sorted based on a 2N DNA content. For each timepoint, Hi-C experiments performed on independent clonal cell lines for NIPBL (NIPBL-A2 and NIPBL-D7) and RAD21 (RAD21-B1 and RAD21-B3) were treated as replicates, with the resulting datasets merged for plotting. (**E**) Example heatmaps showing the consequences of NIPBL (top) and RAD21 (bottom) depletion for 4 and 24 hours at the same locus. Beneath the NIPBL heatmaps are scale factor-normalized RAD21 ChIP-seq tracks for the corresponding treatment in NIPBL-D7 cells. An example chromatin loop is labeled with an arrow in the heatmaps, while the anchors of this loop are marked a grey box in the ChIP-seq tracks.

Given that NIPBL is described as a cohesin loading factor, we performed chromatin fractionation and ChIP-seq in asynchronous cells treated with DMSO or dTAG^V^-1 for 4 or 24 hours to characterize the consequences of both short- and long-term depletion on chromatin association of cohesin. By chromatin fractionation, we observed only a modest reduction in chromatin-associated cohesin, particularly for the NIPBL-D7 cell line (**Figure S2A**). This is despite undetectable NIPBL in the chromatin fraction with dTAG^V^-1 treatment (**Figure S2A**). In comparison, ChIP-seq indicated that RAD21, a core subunit of cohesin, was progressively lost from its binding sites with depletion of NIPBL (**Figure S2B**). Even though the majority of RAD21 peaks were not significantly lost until NIPBL was depleted for 24 hours, almost all peaks showed a degree of reduction at both 4 and 24 hours (**Figure S2B**). Retained and diminished peaks had similar enrichment of HA-NIPBL, but diminished peaks had weaker RAD21 binding in the control DMSO-treated condition (**Figure 1B, Figure S2C**). The observed persistence of RAD21 on chromatin by fractionation coupled with the loss of RAD21 binding at specific sites observed by ChIP-seq suggest that cohesin is able to non-specifically associate with chromatin in the absence of NIPBL. Despite residual chromatin-associated RAD21 with NIPBL depletion, we observed a strong G1 cell cycle arrest with NIBPL depletion, but not with dTAG^V^-1 treatment of parental hTERT RPE-1 cells (**Figure 1C, Figure S2D-F**). In contrast, RAD21 depletion resulted in a G1 cell cycle arrest and an accumulation of 4N cells (**Figure 1C, Figure S2D-F**). Cohesin is more stable on chromatin during S/G2 compared to G1 (*10*), suggesting that a subset of residual chromatin-bound RAD21 with 4-hour NIPBL depletion is derived from S/G2/M cells that have already undergone NIPBL-dependent, stable cohesin loading. In contrast, at 24 hours there are few cells outside of G1 with NIPBL depletion, suggesting that the residual RAD21 is not the product of cohesive cohesin binding to chromatin. Based on these observations, we conclude that RAD21 is capable, albeit less efficient, at binding chromatin in the absence of NIPBL.

### NIPBL-dependence defines distinct classes of chromatin loops

In order to understand the impact of the loss of NIPBL-mediated loop extrusion, we first defined cohesin-dependent genome organization using our RAD21-FKBP12^F36V^ cell line. Given our observation that RAD21-depleted hTERT RPE-1 cells undergo a bimodal cell cycle arrest, with a mix of 2N and 4N cells (**Figure 1C, Figure S2D-F**), we performed the initial steps of Hi-C in these cells before sorting 2N cells based on propidium iodide staining (**Figure S3A**). Comparing the impact of RAD21 or NIPBL depletion on 3D genome organization, the consequences of NIPBL depletion were less severe than with depletion of RAD21 (**Figure 1D, Figure S3B**), consistent with our observation that there is residual RAD21 on chromatin with NIPBL depletion. In contrast with the immediate but persistent consequences of RAD21 depletion, the progressive loss of RAD21 from chromatin with NIPBL depletion is reflected in the progressive reduction in local chromatin contacts (**Figure 1D**). At individual loci, the progressive disruption to RAD21 binding with NIPBL depletion is evident, but did not show an obvious correlation with the rate at which 3D genome contacts are perturbed (**Figure 1E, Figure S3C**).

Given our observation that the consequences of NIPBL depletion were less severe than those with RAD21 depletion and that the extent of these consequences was locus-dependent, we called chromatin loops to more directly address these differences. From control DMSO-treated, FKBP12^F36V^-NIPBL cells, we called 18,825 loops. As it has previously been determined that not all chromatin loops require cohesin (*24*), we classified 16,860 of these loops as cohesin-dependent based on a fold-change greater than two with RAD21 depletion (**Figure S4A**). While cohesin-dependent loops were also strongly perturbed by NIPBL depletion, there was residual signal (**Figure S4B**), consistent with our observations in **Figure 1D**. Indeed, only 42% of cohesin-dependent chromatin loops experienced a fold-change greater than two with NIPBL depletion (**Figure S4C**), highlighting that not all chromatin loops are equally dependent on NIPBL. Based on the loops classified as cohesin-dependent (**Figure 2A, Figure S4A**), we performed unsupervised K-means clustering to distinguish cohesin-dependent chromatin loops based on their sensitivity to NIPBL loss (**Figure 2B**). Overall, the clusters showed a variable loss of loop strength in response to NIPBL depletion, leading to our naming convention (from minimum-to maximum-dependency on NIPBL; **Figure 2B**). Most clusters showed a similar loss of loop strength at both 4 and 24 hours, with the primary exception being a “mixed-dependency” cluster that showed a slight reduction in loop strength at 4 hours and further reduction with 24 hours of NIPBL depletion (**Figure 2B, C**). In contrast, the mixed-dependency cluster was strongly perturbed with 4 and 24 hours of RAD21 depletion (**Figure 2C**), confirming that these chromatin loops are cohesin-dependent.

**Fig. 2:**
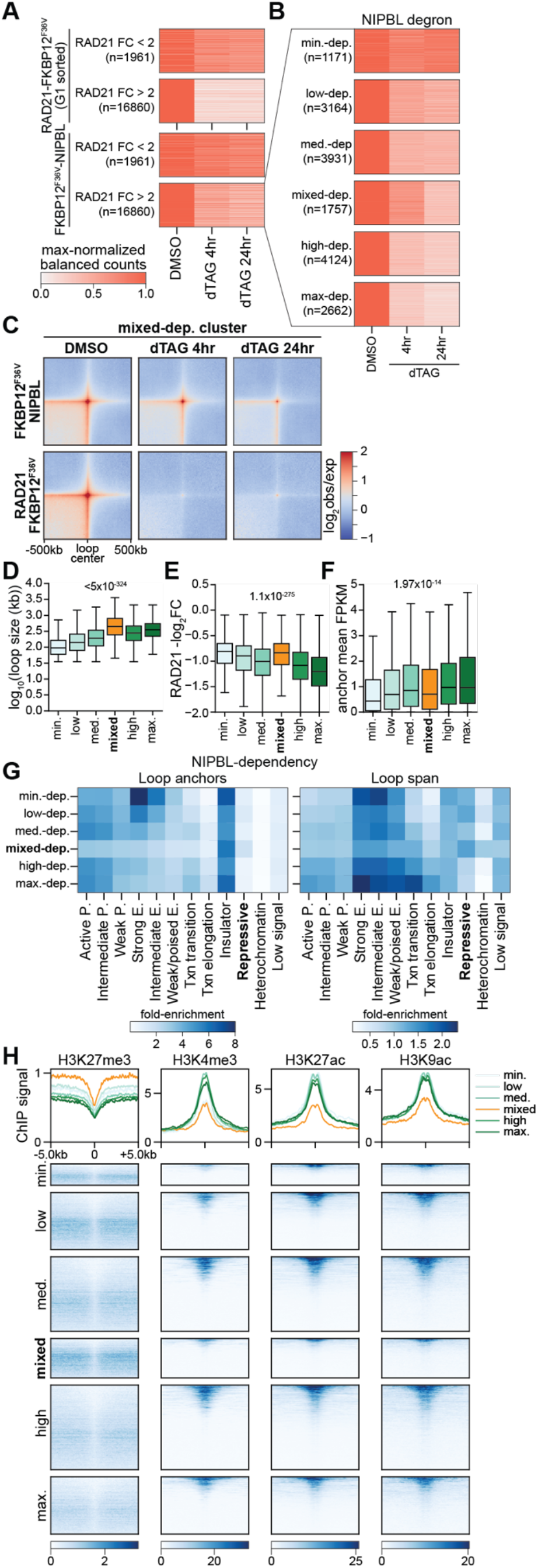
A subset of large chromatin loops, anchored in repressive chromatin, are stable with NIPBL depletion. (**A**) Heatmap showing max-normalized balanced counts of chromatin loops that we classified as RAD21-independent or RAD21-dependent based on a two-fold decrease in chromatin loop strength (balanced counts) in response to RAD21 depletion (top). Also shown are the consequences of NIPBL-depletion on the strength of RAD21-dependent and -independent chromatin loops (bottom). (**B**) Heatmap showing max-normalized balanced counts of K-means clustering of RAD21-dependent chromatin loops from asynchronous cells depleted of NIPBL. Cluster names refer to NIPBL-dependency; minimum-dependency (“min.-dep.”), low-dependency (“low-dep.”), medium-dependency (“med.-dep.”), mixed-dependency (“mixed-dep.”), high-dependency (“high-dep.”), and maximum-dependency (“max.-dep”). (**C**) Observed/expected pile-up analysis at a 10 kb resolution of the mixed-dependency cluster in cells depleted of NIPBL (top) or RAD21 (bottom). (**D**) Log-transformed sizes of chromatin loops from each of the clusters. P-value shown is from a Kruskal-Wallis test. (**E**) RAD21 differential binding at loop anchors in the different clusters. For each chromatin loop anchor, we identified the RAD21 peak that had the smallest -log_2_(fold-change) and found the mean of the two anchors for a given loop. Chromatin loops with no intersecting RAD21 peaks were excluded, but we kept chromatin loops with only one anchor intersecting a RAD21 peak. P-value shown is from a Kruskal-Wallis test. (**F**) Mean FPKM of nascent transcripts whose promoter-proximal regions (TSS -1 kb/+0.5 kb) intersect chromatin loop anchors from the different clusters. Nascent transcripts levels are from SLAM-seq performed on DMSO-treated NIPBL-D7 cells. P-value shown is from a Kruskal-Wallis test. (**G**) Enrichment of ChromHMM chromatin states in cluster loop anchors and loop spans. For loop anchors, both anchors were included in analysis. P = promoter; E = enhancer; txn = transcriptional. Fold-enrichment is calculated by ChromHMM based on the proportions of bases in the chromatin loop anchors/spans that fall into each state compared to the proportion of bases in the whole genome that fall into that state. (**H**) RPKM-normalized ChIP signal for example histone marks used in identifying ChromHMM chromatin states centered at RAD21 ChIP-seq peaks that overlap anchors of chromatin loops in each of the clusters.

We next looked at the features of each cluster to understand why they might be differentially dependent on NIPBL. Chromatin loop size was a clear predictor of dependence on NIPBL (**Figure 2D**), with larger chromatin loops more sensitive to loss of NIPBL. The mixed-dependency cluster was an outlier in this pattern, having the largest median loop size of 450 kb (**Figure 2D**). Based on previously employed metrics for classifying chromatin loops by their relationship with cohesin/CTCF and *cis* regulatory elements (*18*), we found that structural loops (cohesin/CTCF at both anchors) were most common for each cluster (**Figure S4D**; ranging from 38.9% for low-dep. to 53.6% for mixed-dep.). However, less NIPBL-dependent clusters were more likely to be classified as *cis* regulatory element loops (enhancer/promoter at both anchors; **Figure S4D**; ranging from 9.7% for max.-dep. to 25.2% for min.-dep.). For the mixed-dependency cluster, the majority of chromatin loops were structural, with only a minority being *cis* regulatory element loops (**Figure S4D**).

Given our earlier findings that chromatin binding of RAD21 is not lost uniformly across the genome (**Figure S2B**), we tested whether dependence of chromatin loops on NIPBL is related to the strength of RAD21 perturbation. We found that the majority of the mixed-dependency cluster have anchors associated with non-significantly changed RAD21 ChIP-seq peaks (**Figure S4E**). Interestingly, while the clusters showed an inverse relationship between NIPBL-dependence and called RAD21 peaks (less dependent = fewer peaks), the peaks that are associated with less-dependent clusters are more resistant to NIPBL depletion (**Figure S4E**). However, given that the less dependent clusters are still classified as cohesin-dependent, this suggests cohesin is required for the formation of these chromatin loops, but they may then be maintained for short periods of time independent of cohesin. While all chromatin loop anchor-associated RAD21 ChIP-seq peaks showed a level of loss with NIPBL depletion, the degree by which RAD21 is lost was strongly correlated with NIPBL dependency (**Figure 2E**). Consistent with the persistence of the mixed-dependency cluster, the reduction in RAD21 chromatin binding that occurs at its loop anchors was similar to the minimum-dependency cluster (**Figure 2E**).

Subunit composition and post-translational modification have both previously been suggested to influence the size and stability of chromatin loops (*25, 26*). Specifically, STAG1 and STAG2 are mutually-exclusive, core cohesin subunits, with STAG1 thought to support longer and STAG2 shorter chromatin loops (*25*). Consistent with this and our observation that longer chromatin loops are more dependent on NIPBL, we found that STAG1 had higher levels at the more dependent clusters, while STAG2 had higher levels at the less dependent clusters (**Figure S4F**). SMC3 acetylation has been found to enhance the residence time of cohesin on chromatin (*25*). Given this, it is perhaps unsurprising that the clusters of chromatin loops that are least sensitive to NIPBL-depletion in the short-term tended to have higher levels of SMC3ac (**Figure S4F**). Furthermore, the levels of HA-NIPBL were lowest and the levels of PDS5A highest at the mixed-dependency cluster (**Figure S4F**), consistent with the evidence suggesting that NIPBL and PDS5A compete with one-another for binding to cohesin (*13, 27*). Lastly, the cohesin unloading factor WAPL showed only a mild relationship with the different clusters (**Figure S4F**), suggesting that any differential stability of RAD21 and chromatin contacts was not necessarily driven by altered WAPL distribution. Given that the mixed-dependency cluster shared attributes with clusters that are much more dependent on NIPBL, including size, strength, and classification as structural loops, the persistence of these chromatin loops with NIPBL depletion is surprising. However, from these attributes, it is not immediately clear as to why these chromatin loops may be retained with NIPBL depletion.

### Chromatin loop persistence is predicted by chromatin state

In analyzing attributes associated with NIPBL dependency of chromatin loops, we observed that genes associated with the loop anchors of each cluster showed distinct levels of nascent transcription (**Figure 2F**). This raises the possibility that local chromatin states may influence chromatin loop stability. To address this, we used ChromHMM to classify chromatin states (**Figure S4G**). Consistent with our focus on cohesin-dependent chromatin loops for clustering, all clusters were strongly enriched for insulator chromatin states (**Figure 2G**), which are associated with CTCF (**Figure S4G**). Enhancer chromatin states were strongly associated with the anchors and span of loops that are least dependent on NIPBL (min.- and low-dependency; **Figure 2G**), indicating that while at least some enhancer-associated loops require the core cohesin complex, they can persist in the absence of NIPBL. Clusters that were more dependent on NIPBL (high- and max.-dependency) tended to have promoter chromatin states at their anchors, but enhancer states within their loops (**Figure 2G**). In contrast to both the less- and more-dependent clusters, the anchors of the mixed-dependency cluster showed low enrichment with promoter- and enhancer-associated chromatin states (**Figure 2G**), and the histone marks that are prominent in these states (**Figure 2H, S4H**). Instead, the anchors and span of the mixed-dependency cluster showed enrichment in repressive chromatin (**Figure 2G**), which is identified based on the presence of the H3K27me3 mark (**Figure S4G**). The span of chromatin loops in the mixed-dependency cluster was also slightly enriched for heterochromatin (**Figure 2G**). The enrichment of repressive chromatin is consistent with the comparatively low levels of nascent transcripts from associated genes in the mixed-dependency cluster (**Figure 2F**), with this cluster also linked to a disproportionate number of HOX genes (**Figure S4H**). We confirmed enrichment of repressive H3K27me3 marks and scarcity of active marks by looking at the relationship of these with RAD21 ChIP-seq peaks in each of the clusters. We found that, while RAD21 peaks themselves are devoid of H3K27me3 signal, the mixed-dependency cluster shows particularly strong signal flanking the peak (**Figure 2H**). We conclude that the chromatin loops in the mixed-dependency cluster are unique cohesin-dependent chromatin loops, primarily due to their size, association with repressive chromatin, and ability to be maintained in the absence of NIPBL.

### Persistent loops require NIPBL for establishment post-mitosis

Our analysis of 3D genome organization after NIPBL depletion indicated that a subset of RAD21-dependent chromatin loops do not require NIPBL for their ongoing maintenance. While we predicated this analysis on these loops being cohesin dependent, it is unclear whether these loops require NIPBL for their initial establishment or can form through NIPBL independent mechanisms. To address the contributions of NIPBL to the formation of 3D genome structures, we utilized the intrinsic genome-wide reorganization of 3D structures that occurs during mitotic exit, driven by the shift from condensin-to cohesin-mediated structures (*18, 28*). We synchronized NIPBL-D7 cells to mitosis using a double thymidine block and nocodazole, treating cells with DMSO or dTAG^V^-1 simultaneous with nocodazole. Cells were then released from the nocodazole-mediated arrest and cell cycle progression tracked using propidium iodide staining. We first confirmed that depletion of NIPBL from mitotic cells had no major consequences on their ability to enter G1 (**Figure S5A**), consistent with our earlier finding that NIPBL becomes dispensable for cell cycle progression by the completion of G1 (**Figure 1C**). We next evaluated the time window in which we could perform experiments without the G1 cell cycle arrest resulting from NIPBL depletion creating divergence between DMSO- and dTAG^V^-1-treated cells. With this, we found that 8 hours post-release from nocodazole both DMSO- and dTAG^V^-1-treated cells were in G1 (**Figure S5B**). However, by 16 hours post-release, a subset of the DMSO-treated cells has progressed out of G1, while dTAG^V^-1-treated cells remained (**Figure S5C**). Lastly, we verified that HA-NIPBL was being successfully depleted with dTAG^V^-1 treatment of mitotic cells (**Figure S5D**).

To determine the requirement of NIPBL for post-mitotic genome reorganization, we performed Hi-C in NIPBL-D7 cells at 30-minute intervals following release from nocodazole with DMSO-or dTAG^V^-1 treatment (**Figure S6A**). We found that significant structural reorganization occurred with both DMSO and dTAG^V^-1 treatment during mitotic exit, but there was clear disruption to chromatin loop formation in the dTAG^V^-1-treated condition that first became apparent at the 90 minute timepoint (arrow; **Figure S6B**). Indeed, at cohesin-dependent chromatin loops (from **Figure S4A**), the perturbation that occurred with NIPBL depletion during mitotic exit was distinctly stronger at t=120min than with depletion from asynchronous cells (**Figure S6C**). While we did observe a minor strengthening of cohesin-dependent chromatin loops between 120 minutes and 8 hours post-release from nocodazole in both the DMSO- and dTAG^V^-1-treated conditions, the effects of NIPBL depletion during mitotic exit remained more severe at 8 hours than asynchronous depletion (**Figure S6C**). These findings broadly suggest that NIPBL is more critical to the establishment of chromatin loops than their maintenance.

Differentiating the role of NIPBL in chromatin loop maintenance versus formation was of particular interest for chromatin loops that we identified as the “mixed-dependency” cluster, given that these loops share many attributes with clusters that were highly dependent on NIPBL, particularly their size, enrichment in the cohesin subunit STAG1, and relationship with CTCF and cohesin. Despite these attributes, we found that these chromatin loops were comparatively resistant to NIPBL depletion from asynchronous cells at short timescales (**Figure 2C**), possibly due to their association with repressive chromatin (**Figure 2G**) and SMC3 acetylation (**Figure S4F**). Given that NIPBL depletion from cells during mitotic exit caused stronger changes to genome organization compared to depletion from asynchronous cells (**Figure S6C**), we reasoned that the consequences on the mixed-dependency cluster may also be more severe. We found that, across all clusters, NIPBL depletion during the M-G1 transition resulted in weaker chromatin loops than with asynchronous depletion (**Figure 3A**). The enhancer-associated clusters of chromatin loops that we found to be less dependent on NIPBL for their maintenance (min.-dep to med.-dep) were perturbed with NIPBL depletion both 120 minutes and 8 hours post-release from nocodazole (**Figure 3A**). However, there was a notable increase in the strength of these loops by 8 hours after mitotic exit, suggesting that they can partially reform with sufficient time in an NIPBL independent manner. For the mixed-dependency cluster, we observed a substantial loss of loop strength with NIPBL depletion at both 120 minutes and 8 hours post-release (**Figure 3A**). By looking at individual loci within the mixed-dependency cluster, we noted persistence of chromatin loops only with NIPBL depletion from asynchronous cells, but not with depletion during mitotic exit (**Figure 3B**). At these loci, the effect of NIPBL depletion on the strength of RAD21 peaks varied greatly. At the example loci on chromosomes 1 and 2, RAD21 at the anchors is reduced by at least 60%, whereas the upstream chromosome 7 anchor showed minimal perturbation in bound RAD21 (**Figure 3B**). Furthermore, as suggested by the chromatin states from ChromHMM, the anchors of these loops were enriched with H3K27me3 (**Figure 3B**). Our finding that all clusters were more dependent on NIPBL for their establishment may point to a role of NIPBL in facilitating the formation of more stable, higher-order genome structures that provide continued support to cohesin dynamics even when NIPBL is depleted. However, the ability of chromatin loops in the mixed-dependency cluster to persist when NIPBL is depleted from asynchronous cells, but not with depletion during mitotic exit, suggests that this cluster may represent a unique set of chromatin loops that are particularly long-lived.

**Fig. 3:**
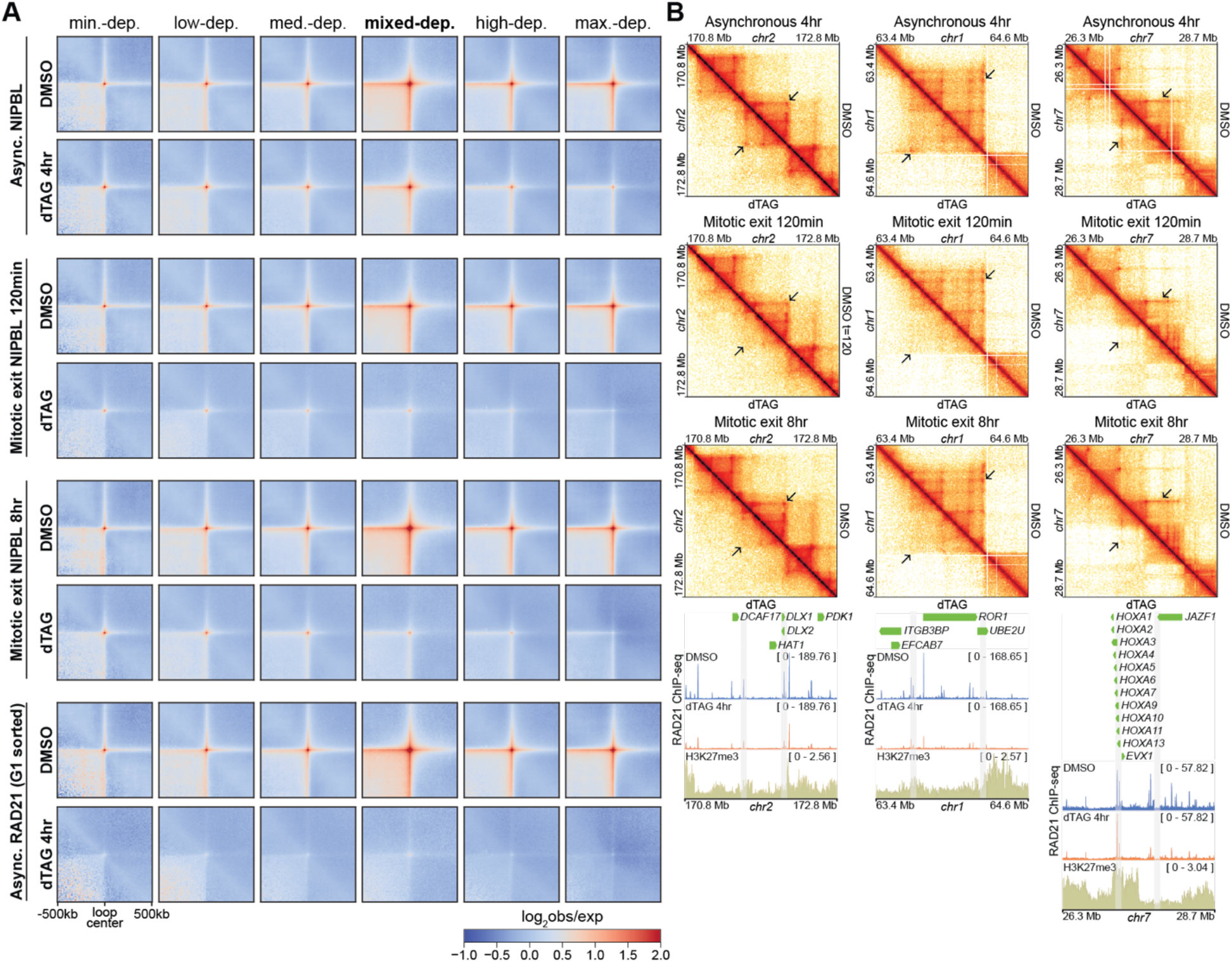
Persistent chromatin loops in the absence of NIPBL. (**A**) Observed/expected pile-up analysis of Hi-C data at a 10 kb resolution of chromatin loops falling into each of the clusters form Figure 2B. (**B**) Example chromatin loops that fall into the mixed-dependency cluster, with Hi-C heatmaps from NIPBL-depletion from asynchronous cells and during mitotic exit, both at early (t=120min) and late (t=8hr) timepoints. Below each locus are a selection of genes that fall into that region, scale factor-normalized BigWig tracks from RAD21 ChIP-seq in asynchronous NIPBL-D7 cells treated with DMSO or dTAG^V^-1 for 4 hours, and an RPKM-normalized BigWig track from H3K27me3 ChIP-seq in hTERT RPE-1 cells. The chromatin loop of interest is marked by an arrow, while its anchor regions are marked by grey bars in the tracks below.

### NIPBL is necessary for activation of lineage-defining genes

Cohesin is reported to preferentially promote gene activation (*29*–*31*). The most prominent instance of gene activation that occurs intrinsically in cells is during mitotic exit (*19*). Whether the shift in 3D genome organization that also occurs during mitotic exit is required for the activation of gene expression is currently unknown. Studying this relationship is made challenging by the role of cohesin in sister chromatid cohesion. However, given our determination that NIPBL can be depleted from cells during mitotic exit without disrupting progression to G1 (**Figure S5A**), we reasoned that our NIPBL-D7 cell line would be an excellent tool to determine if post-mitotic genome organization was a necessary pre-requisite for accurate post-mitotic transcription activation.

We established two approaches to assess the consequences of NIPBL depletion on transcription activation during mitotic exit by SLAM-seq (**Figure S7A**). These approaches were both based on a double thymidine block and nocodazole synchronization but differed in the length of the 4-thiouridine (4sU) labeling period for the identification of nascent transcripts by SLAM-seq (*32*). We reasoned that the longer labeling period (Mitotic exit approach B; **Figure S7A**) would be most suitable for differential analysis, while the shorter labeling periods (Mitotic exit approach A; **Figure S7A**) would allow us to understand the kinetics of nascent transcription, and how this is affected by NIPBL depletion.

We found that depletion of NIPBL during mitotic exit resulted in the differential expression of 549 genes, the vast majority of which were decreased in the absence of NIPBL (**Figure 4A, B**). By gene ontology analysis, genes that failed to activate with NIBPL depletion showed a significant enrichment in processes related to cell behavior, including cell migration and shape (**Figure 4C**). Similarly, gene set enrichment analysis (GSEA) showed a strong negative correlation with genes defining the epithelial to mesenchymal transition (EMT) and those upregulated with KRAS signaling (**Figure 4D**). Indeed, hTERT RPE-1 cells have been described as being dedifferentiated and fibroblast-like, based both on their proteome and morphology, and in comparison to primary epithelial pigment cells (*33, 34*). Furthermore, consistent with our finding that NIPBL is necessary for transcription activation of lineage-defining gene sets, we observe that prolonged depletion of NIPBL results in a shift from a fibroblastic to a more epithelial-like morphology (NIPBL-D7: **Figure 4E**, NIPBL-A2: **Figure S7B**). Taken together these results indicate that NIPBL plays a key role in establishing the expression of gene regulatory programs that are critical for maintaining cell identity.

**Fig. 4:**
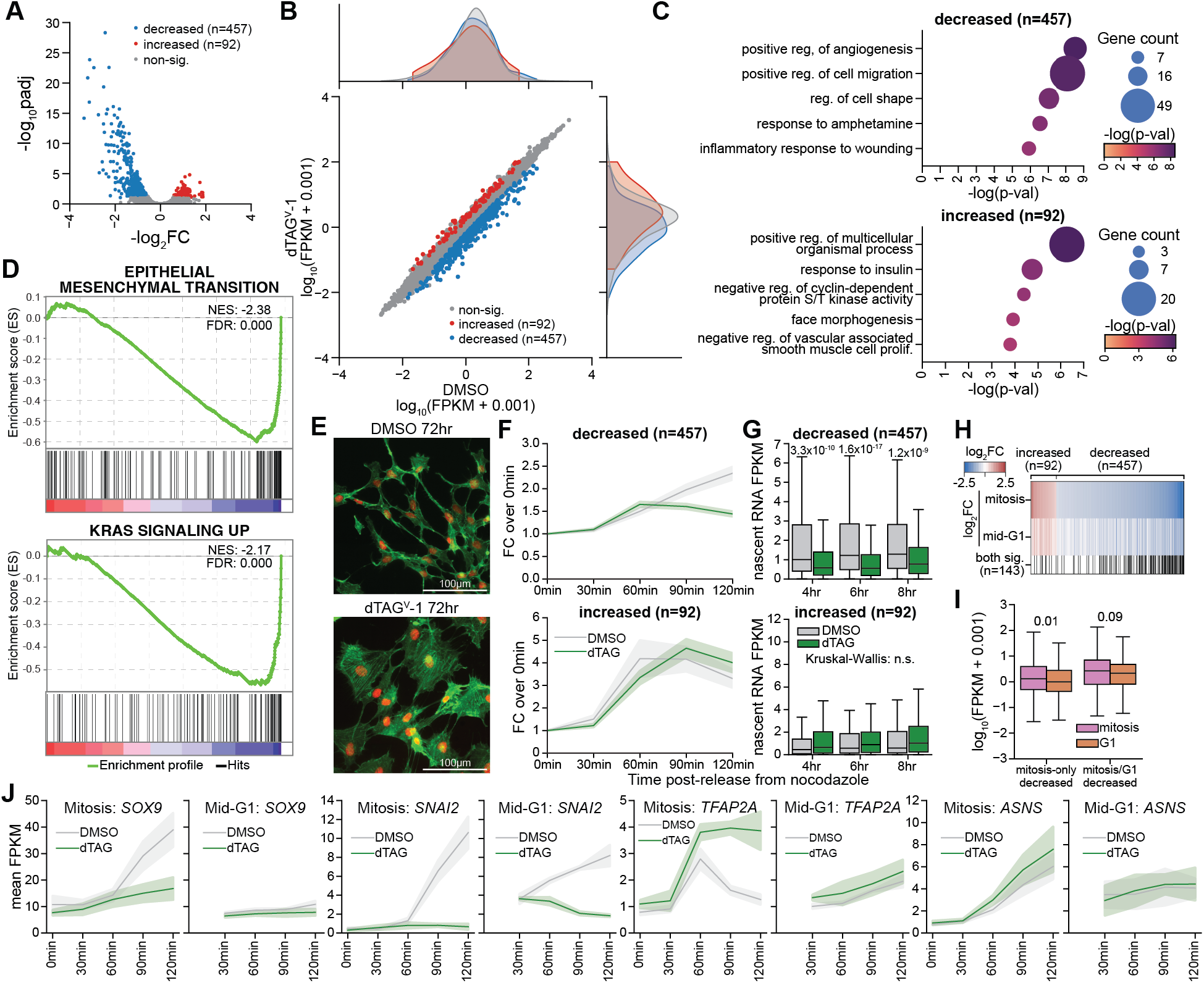
NIPBL facilitates the activation of lineage-defining genes during mitotic exit. (**A**) Volcano plot from differential expression analysis of nascent transcripts between DMSO- and dTAG^V^-1-treated NIPBL-D7 cells based on a 120 min 4sU labeling period during mitotic exit. Significantly changed (|log_2_FC|>0.585 and padj<0.1) nascent transcripts are highlighted. Differential analysis was performed based on 3-4 independent replicates. (**B**) Scatter plot showing the log_10_-transformed FPKM values for nascent transcripts in the DMSO- and dTAG^V^-1-treated NIPBL-D7 cells following a 120 min 4sU labeling period during mitotic exit. Significantly changed transcripts (|log_2_FC|>0.585 and padj<0.1) are highlighted. A kernel density estimate (KDE) plot is shown on each axis to represent the distribution of the datapoints on that axis. To each FPKM value, 0.001 was added prior to log-transformation to allow for plotting of zero FPKM values. (**C**) Biological process gene ontology analysis was performed on the significantly decreased (top) and increased (bottom) nascent transcripts between DMSO and dTAG^V^-1 following a 120 min 4sU labeling period during mitotic exit. The top 5 categories are shown for each analysis, with the p-values derived using the ‘elim’ algorithm from TopGO. (**D**) Gene set enrichment analysis (GSEA) of genes ranked by log_2_FC from differential gene expression analysis of nascent transcripts with 120 min 4sU labeling period. Two “Hallmark” gene sets are shown. (**E**) Morphology of NIPBL-D7 cells treated with DMSO or dTAG^V^-1 for 72 hours. Cells were stained with phalloidin (actin; green) and DAPI (DNA; red). (**F**) Line plots showing the fold-change (FC) in FPKM over t=0min for each timepoint for NIPBL-D7 cells treated with DMSO or dTAG ^V^-1 during mitotic exit (approach A, Figure S7A). Significantly decreased (top) and increased (bottom) genes identified from differential analysis of nascent transcripts with 120 min 4sU labeling period (approach B, Figure S7A) are shown. (**G**) Boxplots showing the FPKM values for nascent transcripts from 15min 4sU labeling period at late timepoints following treatment of NIPBL-D7 cells with DMSO or dTAG ^V^-1 during mitotic exit (approach A, Figure S7A). Significantly decreased (top) and increased (bottom) genes identified from differential analysis of nascent transcripts with 120 min 4sU labeling period (approach B, Figure S7A) are shown. Kruskal-Wallis tests were performed first, before pairwise comparisons were made using two-sided Wilcoxon rank-sum tests. (**H**) Comparison of differential analysis of nascent transcripts with a 120 min 4sU labeling period performed with DMSO-or dTAG ^V^-1-treatment administered during mitotic exit (“mitosis”; mitotic exit approach A, Figure S7A) or 4 hours post-release from nocodazole (“mid-G1”; mid-G1 approach B, Figure S7C). Shown are all significantly changed genes during mitotic exit, ranked by mitosis log_2_FC. Nascent transcripts that also met the criteria for significance (|log_2_FC|>0.585 and padj<0.1) with the mid-G1 approach are denoted with a black line. (**I**) Log-transformed FPKM of nascent transcripts from DMSO-treated samples for genes that are significantly changed with NIPBL depletion either only during mitotic exit or both during mitotic exit and G1. P-values shown are from two-sided Wilcoxon rank-sum tests. (**J**) Example line plots showing the mean FPKM and standard error for nascent transcripts from 15min 4sU labeling period for NIPBL-D7 cells treated with DMSO or dTAG ^V^-1 during mitotic exit (“mitosis”; mitotic exit approach B, Figure S7A) or 4 hours post-release from nocodazole (“mid-G1”; mid-G1 approach A, Figure S7C). *SOX9* is an example of a gene differentially expressed only during mitotic exit, *SNAI2* is differentially expressed during both mitotic exit and mid-G1, and *ASNS* is non-significant during both mitotic exit and mid-G1. For “mitosis”, the x-axis is time post-release from nocodazole, while for “mid-G1”, the x-axis is time post-addition of DMSO/dTAG^V^-1. Additional examples can be found in Figure S7G.

Based on the reported “waves” of transcriptional activation that occur during the M-G1 transition (*19*), we used the mitotic exit approach A (**Figure S7A**) to understand the behavior of differentially expressed genes during this period. Genes that were significantly decreased in the dTAG^V^-1 condition showed a progressive increase in transcript levels with DMSO treatment, beginning at 30-60 minutes post-release from nocodazole (**Figure 4F; top**), at which point cells are transitioning from mitosis to G1 (**Figure S5A**). NIPBL depletion did not blunt this initial gene activation, but continued activation from 60 minutes was weakened (**Figure 4F; top**). In contrast to the significantly decreased genes, genes that were increased with NIPBL depletion during mitotic exit spiked in activation at 60-90 minutes in the DMSO condition then trended downwards (**Figure 4F; bottom**). In the dTAG^V^-1-treated condition, these genes showed a stronger spike in activation (**Figure 4F; bottom**). We found that these perturbations in nascent transcript levels were not a matter of delayed activation in the absence of NIPBL, given that longer timepoints post-release from nocodazole showed little-to-no recovery of significantly decreased genes (**Figure 4G; top**). However, the differences in nascent transcripts that were increased during mitotic exit between DMSO- and dTAG^V^-1-treated samples became non-significant at longer timepoints (**Figure 4G; bottom**), indicating that their heightened expression caused by NIPBL is relatively transient. Overall, these findings indicate that NIPBL is required for complete activation of lineage-defining genes during mitotic exit, and this activation is necessary for the normal expression of these genes during G1.

### Perturbations in nascent transcripts are stronger during mitotic exit than in G1

We saw the cell cycle as a unique opportunity to compare the requirement of NIPBL in activating and subsequently maintaining the expression of the same gene set. To achieve this, we established two approaches analogous to those we used to look at transcriptional activation during mitotic exit, but for application in G1 (**Figure S7C**). As in our mitotic exit approaches, cells were synchronized to prometaphase using a double thymidine block and nocodazole, but instead cells were released for 4 hours (approximately the middle of G1) before addition of DMSO or dTAG^V^-1 (**Figure S7C**). We similarly added 4sU for either 15 minutes prior to harvesting (mid-G1 approach A) or over the course of DMSO/dTAG^V^-1 treatment (mid-G1 approach B; **Figure S7C**). Using this approach, we observed much fewer significantly changed genes with NIPBL depletion (**Figure S7D, E**), but retained enrichment of similar ontologies, including positive regulation of cell migration (**Figure S7F**). Of the 189 nascent transcripts significantly changed with our mid-G1 approach, the majority of these corresponded to nascent transcripts that were significantly changed in our mitotic exit approach (**Figure 4H**). However, this was only a fraction of the genes that were altered by NIPBL depletion during mitotic exit (**Figure 4H**). Furthermore, the extent by which nascent transcripts were affected by NIPBL depletion in mid-G1 was generally weaker than with depletion during mitotic exit (**Figure 4H**). Indeed, genes that were decreased by NIPBL depletion during mitotic exit, but not during G1, showed modestly lower levels of nascent transcripts during G1 compared to mitotic exit, potentially contributing to this distinction (**Figure 4I**). The consequences of NIPBL depletion on nascent transcripts during mitotic exit and G1 is evident when looking at individual example genes (**Figure 4J, Figure S7G**). *SOX9* and *SNAI2* are examples of genes that were both significantly affected by NIPBL depletion during mitotic exit, but only *SNAI2* nascent transcripts were also significantly perturbed by depletion during G1. These genes also serve as excellent examples of the contribution of NIPBL to the activation of lineage defining genes, with *SOX9* and *SNAI2* together capable of inducing EMT of neural epithelial cells (*35*). Given that many chromatin loops connecting enhancers with promoters have been found to persist with depletion of cohesin (*36*) and our observation that chromatin loops are more strongly perturbed with NIPBL depletion during their establishment in mitotic exit than their maintenance in asynchronous cells (**Figure 3**), the degree by which transcription is perturbed during activation versus maintenance may simply be a consequence of the severity by which genome organization is disrupted. Overall, our findings reveal a preferential role of NIPBL in the activation, rather than maintenance, of genes that define cell lineage, with failure to appropriately activate these genes leading to an apparent reversal of EMT.

### A highly connected enhancer neighborhood supports activation of lineage-defining genes

The level of genome reorganization that occurs over the M-G1 transition was extensive, with NIPBL depletion leading to genome-wide failure to appropriately form the local 3D structures that are present in the DMSO-treated condition (**Figure S6B, C**). Despite this, only a fraction of the transcriptome was affected by NIPBL depletion (**Figure 4A**), leading us to ask what makes these genes sensitive. Given our identification of lineage-defining genes as being dependent on NIPBL for activation during mitotic exit, we looked at the relationship of significantly changed genes with enhancers, and how this relationship was influenced by NIPBL depletion. Based on chromatin loops called from the DMSO condition in early G1, we first found that ∼50% of both decreased and increased genes had promoters associated with a chromatin loop anchor (**Figure 5A**). Furthermore, a disproportionate number of these chromatin loops linked these genes to typical- and super-enhancers (**Figure 5A**). While this analysis supports a relationship between enhancer association and sensitivity to NIPBL depletion, loop calling is intrinsically binary and likely underestimates the enhancer-promoter connectivity. To more comprehensively evaluate promoter-enhancer connectivity, we employed an “enhancer neighborhood”, identifying enhancers that fall within 1 Mb either side of the transcription start site (TSS). Both increased and decreased genes exhibited stronger contacts with super- and typical-enhancers compared to non-significantly changed genes (**Figure 5B, C; Figure S8A, B**). For genes significantly decreased in response to NIPBL, the heightened enhancer-promoter connectivity and the dependence of this connectivity on NIPBL was restricted to primarily within 500 kb of promoter-proximal regions (**Figure 5C, Figure S8B**). Given this distance-dependence, we quantified how the enhancer-promoter contact scores were affected by NIPBL depletion specifically for enhancer-promoter pairs separated by no more than 500 kb. We found that, for both typical- and super-enhancers, contacts between decreased genes and their neighborhood enhancers were more strongly perturbed than contacts between non-significant or increased genes and their neighborhood enhancers (**Figure S8C**). The strength and NIPBL-dependence of chromatin contacts between significantly decreased genes and their enhancers was evident when looking at individual loci of genes that were sensitive to NIPBL depletion (**Figure 5D, Figure S9A, B**). These findings indicate that, not only do significantly decreased genes show a higher degree of contact frequency with nearby enhancers, but their ability to come into contact with these enhancers is more sensitive to NIPBL depletion.

**Fig. 5:**
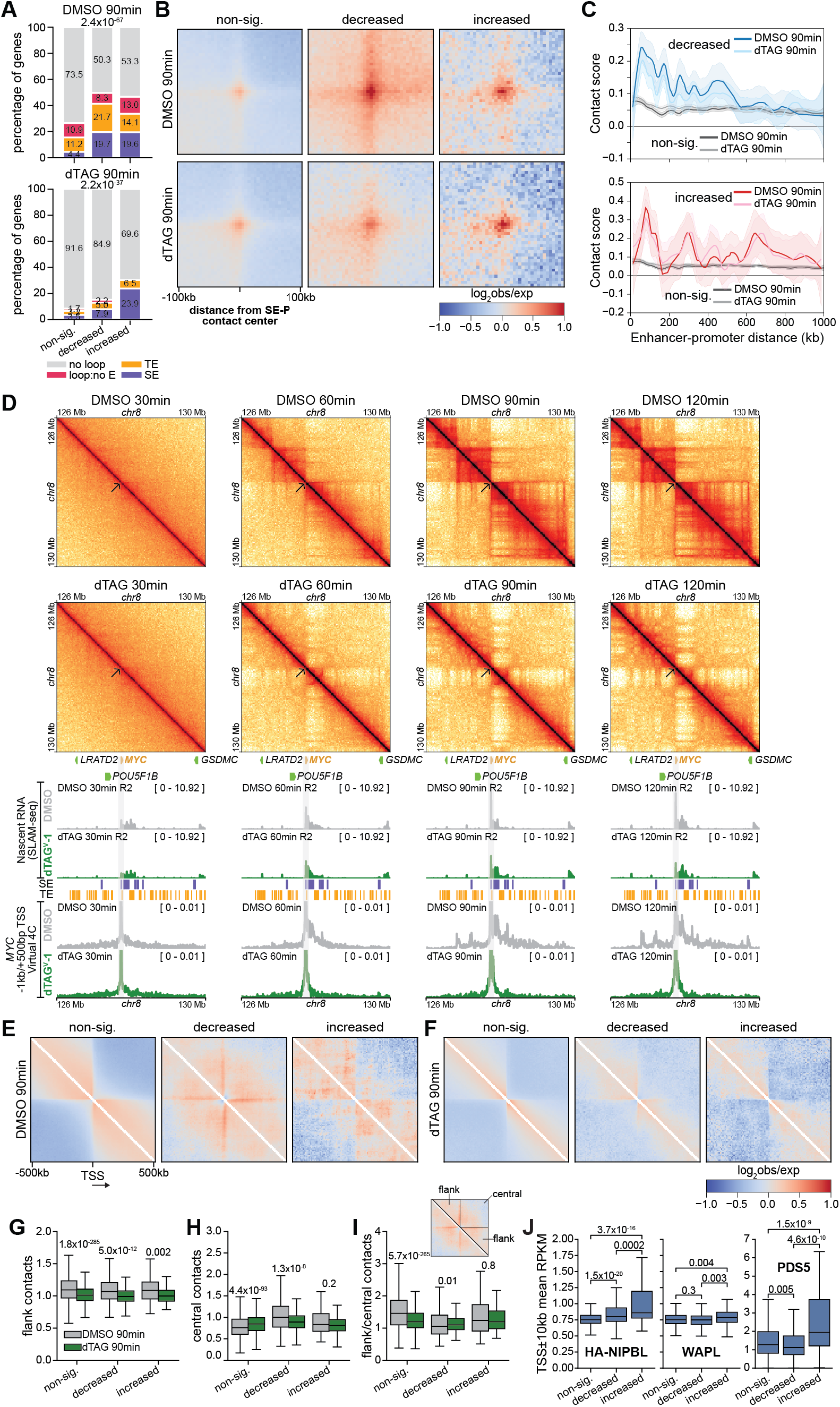
Super-enhancer neighborhood connectivity and local genome organization are associated with sensitivity to NIPBL depletion. (**A**) Stacked bar plot showing the percentage of non-significant, decreased, and significantly increased genes whose promoter-proximal regions (TSS -1 kb/+500 bp) intersect a chromatin loop anchor and whereby the distal loop anchor intersects with a super-enhancer (SE; purple), typical enhancer (TE; yellow), or no enhancer (loop:no E; magenta). Because promoter-proximal regions may intersect with multiple chromatin loop anchors, each gene is represented only once, with chromatin loops for a given gene preferentially selected based on intersection of the distal loop anchor with a super-enhancer, then typical enhancer, then no enhancer. P-values for chromatin loops called from the DMSO- and dTAG^V^-1-treated conditions were determined using a chi-squared test. (**B**) Off-diagonal pile-up analysis of the super-enhancer neighborhood (TSS ± 1Mb) for non-significant, decreased, and increased genes with NIPBL depletion during mitotic exit. Shown is the log-transformed mean observed/expected contact frequency between promoter-proximal regions (TSS -1 kb/+500 bp) and all neighborhood super-enhancers called from H3K27ac ChIP-seq with a TSS exclusion value of 2500 bp and a stitch value of 12500 bp. Enhancers within 10 kb of promoter-proximal regions were excluded. (**C**) Loess-smoothened relationship with 95% confidence interval at a 5 kb resolution between distance and contact frequency between super-enhancers and promoter-proximal regions. The same non-significant contacts are repeated for comparison with the decreased and increased contacts. (**D**) *MYC* is an example of a gene that is significantly decreased in response to NIPBL depletion during mitotic exit. Shown are Hi-C heatmaps (top row: DMSO; bottom row: dTAG), sequencing-depth normalized BigWig tracks of read pairs whereby at least one read contains a T ⟶ C conversion (i.e. nascent reads), and Virtual 4C anchored at *MYC* promoter-proximal regions (TSS -1 kb/+500 bp) for both DMSO and dTAG-treated samples at t=30min, 60min, 90min, and 120min. The marked TE and SE sites are the same across all plots. The arrow on the Hi-C heatmaps marks the *MYC* transcription start site, and the grey bar on the tracks below marks the gene location. (**E & F**) On-diagonal pile-up analysis of Hi-C data at a 10 kb resolution from DMSO-(**E**) and dTAG^V^-1-(**F**) treated cells harvested 90 minutes post-release from nocodazole. Pile-up was centered on transcription start sites (TSSs) of non-significant, decreased and increased genes from Figure 4A, with all genes oriented in the same direction (right-facing). (**G & H**) Quantification of (E) & (F), specifically the mean observed/expected contact frequency for the flanking (**G**) and central (**H**) regions for each TSS. P-values were determined by independent, two-sided t-tests. (**I**) Quantification of (E) & (F), specifically the ratio between observed/expected contacts of regions flanking the TSS (“flank”) and the observed/expected contacts of regions across the TSS (“central”). The inset shows the regions that were quantified, with the mean observed/expected contact frequency for a given TSS taken from the shown regions for plotting. P-values were determined using two-sided Wilcoxon rank-sum tests. (**J**) Mean RPKM of WAPL, PDS5, and HA-NIPBL ChIP-seq at a 20 kb region centered on the TSSs of non-significant, decreased and increased genes. Two-sided Wilcoxon rank-sum tests were performed to give the shown p-values.

### Local chromatin connectivity correlates with sensitivity to NIPBL depletion

Genes have been found to function as barriers to loop extrusion (*37, 38*), giving rise to an insulative effect whereby genomic regions upstream of a TSS tend not to come into close spatial proximity with regions downstream of a TSS. Curiously, we noticed that genes sensitive to NIPBL had a degree of permissiveness to this insulation, in that some contacts were able to form between regions either side of the TSS (**Figure S10A**). To quantify this effect genome-wide, we performed an on-diagonal pile-up of 3D genome contacts centered at the TSS of non-significant and significantly changed genes. With this approach, we found that chromatin contacts between regions up-and down-stream of TSSs were substantially stronger for decreased genes compared to non-significantly changed genes (**Figure 5E**). The contact domains on either-side of the TSS were strongly perturbed by NIPBL depletion for all genes (**Figure 5F, G**), but only with the decreased genes were the cross-TSS contacts between the domains also lost with NIPBL depletion (**Figure 5F, H**). Indeed, the degree of contacts across the TSS is very similar to the contacts up-or down-stream of the TSS, with the ratio of these for decreased genes being close to one and only modestly affected by NIPBL depletion (**Figure 5I**). Curiously, such a contact pattern as we observe at decreased genes has previously been recapitulated globally at TSSs by WAPL depletion (*37*), suggesting that cohesin removal by WAPL may normally limit the processivity of cohesin across the TSS. Given that NIPBL competes with PDS5 for binding to cohesin (*13, 27*), and that association of PDS5 with cohesin is a precursor to cohesin removal by WAPL (*39*), we reasoned that an altered NIPBL-PDS5-WAPL balance at certain loci may facilitate improved cross-TSS processivity of cohesin. We addressed this possibility by quantifying promoter-proximal HA-NIPBL, WAPL, and PDS5, finding that NIPBL was higher and PDS5 lower at significantly decreased genes, whereas levels of WAPL was consistent between non-significant and decreased genes (**Figure 5J**). Given the ability of NIPBL to stimulate the ATPase activity of cohesin (*13*), the imbalance of NIPBL and PDS5 may reflect distinct cohesin dynamics at these loci facilitating stronger interconnectivity of the local 3D genome that is necessary for their activation during mitotic exit.

Enhanced cohesin dynamics at a given locus may also promote the formation of more complex 3D genome structures, such as multi-way contacts between a gene and numerous enhancers. Consistent with this hypothesis and the absence of TSS-centered insulation, we found that both super- and typical-enhancers in the neighborhood of genes sensitive to NIPBL depletion showed strong, NIPBL-dependent engagement with enhancers on the opposite side of the TSS (**Figure S10B**). The cross-TSS connectivity of enhancers was evident at *JARID2*, but was comparatively much weaker at *NECTIN3*, an example of a non-significantly changed gene with NIPBL depletion (**Figure S10C**). Thus, we conclude that NIPBL-mediated loop extrusion regulates gene activity by facilitating a combination of enhancer-promoter and enhancer-enhancer contact across three-dimensional space, possibly by enabling the formation of higher-order structures.

## Discussion

Here, we have characterized the role of cohesin dynamics in regulating 3D genome organization and transcription activation by targeting the cohesin accessory protein NIPBL. Loop extrusion facilitated by NIPBL is more critical to the establishment of chromatin loops upon mitotic exit than their maintenance in asynchronous cells, but generally NIPBL promotes the formation of larger chromatin loops. Despite this relationship, we also identified a subset of large chromatin loops that were particularly persistent when NIPBL is depleted from asynchronous cells, but failed to form when NIPBL is depleted during mitotic exit, suggesting that they may represent “long-lived” chromatin loops. Our capacity to impair cohesin dynamics with NIPBL depletion also gave us the unique opportunity to characterize the differential role of 3D genome organization in facilitating transcription activation in early G1 and steady-state transcription in mid-G1. We found that NIPBL-mediated loop extrusion preferentially activates a core set of lineage-defining genes during mitotic exit, with only a subset of these requiring NIPBL for steady-state expression in G1. Based on identification of this gene set, we determined that sensitivity of gene expression to NIPBL depletion is correlated with disproportionately strong associations with super- and typical-enhancers, and a unique cross-TSS permeability of 3D genome contacts, particularly between enhancers. Overall, our findings provide comprehensive insight into the role of NIPBL in transcription regulation and genome organization across multiple contexts.

The role of NIPBL in regulating loop extrusion and 3D genome organization has not been widely explored *in vivo*. By comparing the consequences of NIPBL and RAD21 depletion in the same cell line, we found that the dependence of cohesin-mediated chromatin loops on NIPBL is strongly related to distance. Specifically, longer chromatin loops are more robustly perturbed by NIPBL depletion than shorter chromatin loops. While this finding is consistent with a role of NIPBL in promoting loop extrusion, a more nuanced perspective is that NIPBL increases the likelihood that cohesin is capable of traversing long distances. The mechanism by which NIPBL achieves this is likely two-fold. First, NIPBL has been found to increase the ATPase activity of cohesin (*13*). Second, NIPBL competes with PDS5 for binding to cohesin, which likely confers a level of resistance to PDS5/WAPL-mediated removal of cohesin from chromatin (*13, 27, 39*). Through these mechanisms, NIPBL may facilitate cohesin processivity to allow for larger chromatin loops to form. Given prior work has found that NIPBL is required for cohesin-mediated loop extrusion *in vitro* (*11, 12*), our findings suggest that other factors may support loop extrusion *in vitro*, and these would be necessary for the formation of shorter-range contacts *in vivo*. While diffusion of cohesin may contribute to the formation of shorter chromatin loops, this is likely severely size-limited (*40*).

By depleting NIPBL from cells during both steady-state and mitotic exit, we have also established the differential role of NIPBL in genome organization “maintenance” versus “formation”. While the 3D genome is highly dynamic, the scale and complexity of reorganization that occurs during the M-G1 transition far exceeds that of cells at steady-state. Indeed, our findings demonstrate that the consequences on genome organization of NIPBL depletion during mitotic exit were more severe than when NIPBL was depleted from asynchronous cells. There are multiple possible mechanisms by which this difference may arise, such as stable compartmentalization of the genome during interphase. However, we also identified a subset of chromatin loops that were disproportionately persistent when NIPBL was depleted from asynchronous cells (the mixed-dependency cluster). We were particularly interested in this chromatin loop set for multiple reasons; they are large in size, RAD21-dependent, anchored in repressive chromatin, retained more RAD21 when NIPBL was depleted, and were highly dependent on NIPBL for formation during mitotic exit. In contrast to the findings that chromatin loops can persist for only 10-20 minutes (*8, 9*), we believe that this set of chromatin loops may represent a distinct type of loop whose longevity is greater than other loops of a similar size. The enrichment of repressive chromatin may shelter cohesin, allowing the chromatin loop to persist for longer periods of time.

Enhancer-promoter connectivity is increasingly viewed as a more complex, higher-order structure than a single enhancer being in close proximity to a single promoter. This complexity may include condensates or hubs involving multiple distal elements (*41*). While Hi-C is mostly ineffective for detecting these complex structures, we did observe a distinct local 3D genome architecture at genes that were sensitive to NIPBL depletion. Specifically, regions upstream of the TSS of sensitive genes were highly connected with regions downstream, whereas up- and down-stream regions were more insulated from one-another at non-sensitive genes. This insulation effect has previously been attributed to the interaction of cohesin with the transcription machinery, and is lost with WAPL depletion (*37*). We posit that cohesin dynamics at sensitive genes are enhanced by higher levels of NIPBL, which may facilitate ongoing loop extrusion at certain genes despite collisions with RNA polymerase. Furthermore, this unique organization may also reflect the formation of higher order 3D genome structures, including simultaneous association of enhancers both up- and down-stream of sensitive genes with their promoter. Indeed, an imaging-based approach, ORCA, has recently been used to demonstrate multi-way contacts by *SOX9* (*42*), which we found to be sensitive to NIPBL depletion during mitotic exit. Similarly, a three-way “kissing” model for cohesin-dependent transcriptional bursting has been found to regulate *Sox2* expression (*43*). Thus, our observation of cross-TSS connectivity may be a consequence of two cohesin complexes converging on a given TSS, resulting in distal elements either side of the TSS coming into a close spatial proximity that we can observe by Hi-C. Our finding that genes sensitive to NIPBL depletion during mitotic exit tend to have such distal connectivity then suggests that this organization is important to gene regulation. The findings presented here are critical steps to understanding multiple aspects of the loop extrusion process and genome organization, providing insights into their form and function both during mitotic exit and at steady state.

## Acknowledgments

We thank Cheng-Zhong Zhang for providing a comprehensive list of SNPs in hTERT RPE-1 cells.

## Author contributions

Conceptualization: TMP and JRD. Formal Analysis: TMP. Funding acquisition: JRD. Investigation – cell line engineering: TMP and FM. Investigation – western blotting: TMP. Investigation – slide preparation and imaging: TMP and MEB. Investigation – propidium iodide staining: TMP and AP. Investigation – mitosis sample collection: TMP and NH. Investigation – ChIP-seq, SLAM-seq, Hi-C: TMP. Methodology: TMP and JRD. Supervision: JRD. Visualization: TMP. Writing – original draft: TMP and JRD. Writing – review & editing: TMP, AP, FM, MEB, NH, and JRD.

## Data and materials availability

Cell lines, plasmids, and other materials are available upon request. Hi-C, SLAM-seq, and ChIP-seq data collected for the purposes of this study are available at the Gene Expression Omnibus (https://www.ncbi.nlm.nih.gov/geo/) with the accession number GSE277721. All datasets generated in this study are presented in **Table S2-4**. Datasets accessed from Sequence Read Archive (SRA) are listed in **Table S5**. Code used for alignment of processing of Hi-C (https://doi.org/10.5281/zenodo.13835562) and SLAM-seq (https://doi.org/10.5281/zenodo.13835597) sequencing data are available from Zenodo.

