## Supplementary Material for "Genome-wide *in vivo* dynamics of cohesin-mediated loop extrusion and its role in transcription activation"

### Materials and Methods

Cell culture

hTERT RPE-1 cells were obtained from ATCC (CRL-4000) and maintained in DMEM/F12 (Thermo Fisher #11330-057) supplemented with 10% FBS (Gibco #A5256701) and 1% penicillin-streptomycin (Gibco #15140122). NIH/3T3 cells were obtained from ATCC (CRL-1658) and maintained in DMEM with 4.5 g/L glucose, L-glutamine and sodium pyruvate (Corning 10-013-CV) supplemented with 10% FBS (Gibco #A5256701) and 1% penicillin-streptomycin (Gibco #15140122).

DMSO (Sigma-Aldrich #D2650) or 500 nM dTAG$^{V}$-1 (Tocris #6914) was added to cells for the indicated periods of time.

Generation of dTAG cell lines

Because *NIPBL* is reported to encode multiple splicing isoforms that vary at the C-terminus of the protein (*44*), we focused on introducing FKBP12$^{F36V}$ to the N-terminus of NIPBL. To achieve this, we manipulated pCRIS-PITChv2-Puro-dTAG (BRD4), a gift from James Bradner and Behnam Nabet (Addgene plasmid #91793; http://n2t.net/addgene:91793; RRID:Addgene_91793) (*20*) to include a mCherry or mGreenLantern reporter in place of PuroR, and replaced *BRD4* homology arms with 200 bp upstream and 811 bp downstream homology arms from *NIPBL*. The pCRIS-PITChv2-mCherry/mGreenLantern-FKBP12$^{F36V}$-NIPBL vectors were further modified by mutation to the PAM sequence by site-directed mutagenesis to prevent off-target cutting by the sgRNA targeting the sequence AGAGTAGTAATGGGGACATG, present towards the end of Exon 2 in *NIPBL*. This sgRNA sequence was introduced into pX333, a gift from Andrea Ventura (Addgene plasmid # 64073 ; http://n2t.net/addgene:64073 ; RRID:Addgene_64073) (*45*).

For RAD21, we focused on creating C-terminal fusion with FKBP12$^{F36V}$ as acute depletion had previously been achieved with the AID system at the C-terminus (*24*, *46*). For this targeting vector, we reorganized the pCRIS-PITChv2 vector to create RAD21-FKPBP$^{F36V}$-2XHA-T2A-mGreenLantern, with 200 bp upstream and 845 bp downstream homology arms for *RAD21*. The sgRNA sequence used to engineer the RAD21 cell line intersected with its stop codon, eliminating any need to adjust the sgRNA target regions in the targeting vectors. The *RAD21* sgRNA, CCTTATATAATATGGAACCT, was cloned into pX330-U6-Chimeric_BB-CBh-hSpCas9, a gift from Feng Zhang (Addgene plasmid #42230; http://n2t.net/addgene:42230; RRID:Addgene_42230) (*47*).

For NIPBL, the targeting and PX330 plasmids were introduced using the Lonza 4D-Nucleofector X with the P3 Primary Cell Kit (Lonza #V4XP-3024), while for RAD21 these were introduced using Takara Xfect Transfection Reagent (#631318). For the NIPBL-D7 and RAD21 degron cell lines, single mGreenLantern positive cells were sorted into individual wells of a 96-well plate. For the NIPBL-A2 degron cell line, mCherry positive cells were first sorted in bulk into a single well of a 6-well plate, but instead these cells were expanded before being subject to a second round of electroporation using the same sgRNA and targeting vector. All sorts were performed into 50% conditioned media using BD Influx Cell Sorters in the Salk Flow Cytometry Core. See **Fig. S1B** for an example gating strategy.

For each of the different degron cell lines, homozygous insertion of the targeting vector payload was initially confirmed by extraction of gDNA by QuickExtract (Biosearch Technologies # QE09050) and PCR using primers outside of the homology arms (see **Table S1** for genotyping primer sequences). Positive clones were expanded, with the genotyping confirmed with gDNA extracted using DNeasy Blood & Tissue (Qiagen #69506), and the same primer pairs in PCR. These PCR products were either directly submitted for Sanger sequencing or introduced into Zero Blunt TOPO vector (Invitrogen #K280020) before Sanger sequencing to confirm accurate insertion of the FKBP12$^{F36V}$ construct. Clones were further validated by western blot, specifically looking at the molecular weight shift of the protein of interest compared to parental hTERT RPE-1 cells, successful depletion of the fusion protein with the addition of dTAG$^{V}$-1, and presence of an HA-tagged species at the appropriate molecular weight.

Whole cell extracts for western blot

Cells were washed in ice-cold Dulbecco's Phosphate-Buffered Saline (DPBS; Gibco #14190144), harvested, and cell pellets resuspended in Kischkel buffer (50 mM Tris pH 8.0, 150 mM NaCl, 5 mM EDTA, 1% Triton X-100) supplemented with 1X cOmplete, EDTA-free Protease Inhibitor Cocktail (PIC; Roche #COEDTAF-RO) and 1 mM PMSF. Cells were sonicated at 25% for 15 seconds using an Active Motif Epishear Probe Sonicator. Debris was removed by centrifugation at 20000 x g for 10 minutes at 4°C. Total protein was quantified using 1X Protein Assay Reagent (BioRad #5000006) against a BSA standard curve. Samples were typically balanced in a new tube to a final concentration of 1 mg/ml using Kischkel buffer and Laemmli buffer (300 mM Tris pH 6.8, 32% glycerol, 8% SDS, bromophenol blue) supplemented with BME, before incubation at 95°C for 5 minutes.

Chromatin fractionation

Cells were subject to chromatin fractionation as described by Mendez and Stillman (*48*). Briefly, cells were washed with DPBS, harvested and resuspended in ice-cold Buffer A (10 mM HEPES pH 7.9, 10 mM KCl, 1.5 mM $MgCl_2$, 0.34 M sucrose, 10% glycerol) supplemented with 1 mM DTT, 1X PIC, and 1 mM PMSF. Triton X-100 was added to a final concentration of 0.1%, and cells were incubated on ice for 8 minutes. The reaction was centrifuged at 1700 x g for 5 minutes at 4°C. The supernatant (fraction S1) was transferred to a new tube, clarified by centrifugation (20000 x g for 10 minutes at 4°C), resulting in the cytosolic fraction (fraction S2). The pelleted nuclei (fraction P1) were gently washed with Buffer A, resuspended in Buffer B (3 mM EDTA, 0.2 mM EGTA) supplemented with 1 mM DTT, 1X PIC, and 1 mM PMSF, and incubated on ice for 30 minutes. The reaction was centrifuged at 1700 x g for 5 minutes at 4°C, with the resulting supernatant (fraction S3; nucleoplasm) transferred to a new tube. The insoluble pellet (fraction P3; chromatin) was gently washed with Buffer B, resuspended in 1X Laemmli Buffer supplemented with BME, and sonicated at 25% for 15 seconds using an Active Motif Epishear Probe Sonicator. Laemmli buffer supplemented with BME was added to 1X to fractions S2 and S3. All samples were incubated at 95°C for 5 minutes.

Western blot

Protein samples in Laemmli buffer were separated out by SDS-polyacrylamide gel electrophoresis (PAGE) using Mini-PROTEAN TGX Pre-Cast Gels (Bio-Rad #4561085) in running buffer (Bio-Rad #1610772). Samples were transferred by wet transfer to PVDF (Bio-Rad Immun-Blot #1620177) using Towbin Transfer Buffer (25 mM Tris, 192 mM glycine, 10% methanol), with the membrane subsequently blocked using 5% non-fat milk in TBS-T (Growcells 10X TBS #MRGF-6690-010L; 0.1% Tween-20). Membrane was incubated overnight at 4°C with primary antibody diluted in 5% non-fat milk in TBS-T and then incubated for 1 hour at room temperature with HRP-conjugated secondary antibody. For visualization, membranes were incubated with ECL substrate (Thermo Scientific SuperSignal West Pico PLUS #34580 and Thermo Scientific SuperSignal West Femto #34095) and imaged using a Bio-Rad ChemiDoc MP.

Antibodies: For western blotting of NIPBL, rabbit polyclonal antibodies from Bethyl (#A301-779A) or Proteintech (#18792-1-AP) were used independently at 1:1000 or combined at 1:1000 each. Other primary antibodies used for western blotting are anti-RAD21 C-terminus (Bethyl #A300-080A; 1:1000), anti-RAD21 N-terminus (Abcam #ab154769; 1:1000), anti-SMC3 (Bethyl #A300-060A; 1:25000), anti-α-Tubulin (Millpore CP06100UG 1:5000), anti-Histone H3 (Cell Signaling #4499, 1:2000), anti-HA (Cell Signaling #3724; 1:1000), and anti-MAU2/SCC4 (Abcam #ab183033; 1:1000). HRP-conjugated secondary antibodies used are goat anti-mouse IgG (Jackson #115-035-071; 1:5000) and goat anti-rabbit IgG (Invitrogen #31463; 1:5000).

Cell cycle synchronization

hTERT RPE-1 cells were synchronized to prometaphase using a double thymidine block and nocodazole as previously described (*49*). Briefly, cells were treated with 2 mM thymidine (Sigma-Aldrich #T1895) for 14 hours, released for 10 hours, then treated again with 2 mM thymidine for 14 hours. Cells were then released for 8 hours before treatment with 20 ng/mL nocodazole (Sigma-Aldrich #M1404) for 2 hours. For experiments aimed at studying post-mitotic transcriptional reactivation, treatments (DMSO or dTAG$^{V}$-1) were initiated simultaneous with nocodazole. Mitotic cells were then collected by shake-off, washed once, before being released into fresh media with or without treatment. For experiments aimed at determining gene expression changes within the G1 phase, cells were released from nocodazole for four hours before addition of DMSO or dTAG$^{V}$-1. For all experiments, cells were harvested at the stated timepoints post-nocodazole release for propidium iodide staining, western blotting, SLAM-seq, or Hi-C.

Cell cycle analysis by propidium iodide staining

Cells treated with DMSO or dTAG$^{V}$-1 were harvested at the stated times, immediately fixed in ice-cold 90% ethanol, and incubated at -20°C for at least 2 hours. Cells were brought to room-temperature and washed 1X with DPBS. Fixed cells were then resuspended in propidium iodide staining buffer (10 µg/ml propidium iodide (Abcam #ab14083), 100 µg/ml RNase (Thermo Scientific #EN0531), 2 mM $MgCl_2$, DPBS) and incubated overnight at 4°C. Flow cytometry analysis was performed using either a BD LSR II or BD FACSymphony A3 in the Salk Flow Cytometry Core. An example of the gating strategy used for these experiments is shown in **Fig. S2**.

Cell imaging

Cells were grown on glass coverslips for 24 hours before being treated with DMSO or dTAG$^{V}$-1 for 72 hours. Coverslips were transferred to a 24-well plate, washed 2X with DPBS, and fixed in 4% methanol-free formaldehyde for 15 minutes at room temperature. Following 2X washes with DPBS, cells were permeabilized with 0.1% Triton X-100 for 15 minutes at room temperature and then washed 1X in DPBS. Cells were then stained using 1X Alexa Fluor 488 Phalloidin (Invitrogen #A12379) in DMSO for 45 minutes at room temperature. DAPI (Thermo Scientific #D1306) was then added to a final concentration of 0.35 ng/µl, incubating for a further 15 minutes at room temperature. Coverslips were washed 2X with DPBS and then mounted to slides using Prolong Diamond Antifade Mountant (Invitrogen #P36965). Slides were then imaged using the Olympus VS-120 Virtual Slide Scanning Microscope.

Chromatin immunoprecipitation for sequencing (ChIP-seq)

Chromatin preparation for ChIP-seq using antibodies against CTCF, RAD21, PDS5A, WAPL, STAG1, and STAG2 was performed by fixing ~10 million cells with 1% methanol-free formaldehyde (Thermo Scientific # 28908) for 10 minutes, then quenching with 125 mM glycine for an additional 10 minutes. For HA-NIPBL ChIP-seq, ~10 million cells were fixed with 2 mM DSG (Santa Cruz #sc-285455) for 45 minutes, washed 3X with DPBS, fixed further with 1% methanol-free formaldehyde for 10 minutes, and quenched with 125 mM glycine for an additional 10 minutes. Cells were washed 2X in DPBS, harvested, flash frozen and stored at -80°C until proceeding. Cells were then lysed for 15 minutes on ice using formaldehyde lysis buffer (FALB; 50 mM HEPES pH 7.5, 140 mM NaCl, 1 mM EDTA, 1% Triton X-100) supplemented with 1% SDS and 1X PIC, and sonicated for 9 minutes (10 seconds on/10 seconds off) at 25% power using an Active Motif Epishear Probe Sonicator. Debris was removed by centrifugation at 21000 x g for 10 minutes at 4°C. For immunoprecipitation, 30 µl Protein A Dynabeads (Invitrogen #10002D) were washed 3X with FALB (no SDS) and blocked using 10 µg of BSA. Antibody and beads were incubated together for 6-8 hours rotating at 4°C, after which the beads were washed 3X in FALB (no SDS). An "input" sample of the prepared chromatin was taken out before the remaining chromatin was diluted out 10-fold with FALB to lower the SDS concentration, and antibody-bound beads were added, incubating overnight rotating at 4°C. Following incubation, beads were washed 1X for 5 minutes rotating at 4°C in each of the following ice-cold buffers: high-salt wash buffer (20 mM Tris pH 8.0, 500 mM NaCl, 2 mM EDTA, 0.1% SDS, 1% Triton X-100), lithium chloride wash buffer (10 mM Tris pH 8.0, 250 mM LiCl, 1 mM EDTA, 1% IGEPAL), and TE wash buffer (10 mM Tris pH 8.0, 1 mM EDTA, 50 mM NaCl). Beads were incubated in elution buffer (50 mM Tris pH 8, 1 mM EDTA, 1% SDS) for 1 hour at 65°C, with the supernatant transferred to a new tube. "Input" and ChIP samples were supplemented with NaCl to 200 mM and 40 µg proteinase K (NEB #P8107S), incubated for 30 minutes at 55°C and then overnight at 65°C. The following day, samples were treated with RNAse A (Thermo Scientific #EN0531) for 2 hours at 37°C, before purification using the Zymo ChIP DNA Clean & Concentrator kit (#D5205).

For experiments that included a 10% spike-in of NIH/3T3 cells (RAD21 ChIP), the NIH/3T3 cells were fixed, then lysed and sonicated using the required buffer and conditions. Chromatin prepared from the NIH/3T3 cells was then combined with the chromatin prepared from the engineered hTERT RPE-1 cells (~10% NIH/3T3 + ~90% hTERT RPE-1 by cell number) prior to dilution and immunoprecipitation.

For CTCF and HA-NIPBL ChIP-seq, "input" and ChIP samples were subject to a dual-sided size selection using AMPure XP beads (Beckman Coulter # A63880) before sequencing libraries were prepared using 2S Plus DNA Library Kit (IDT #10009878). For RAD21, WAPL, PDS5A, STAG1, and STAG2 ChIP-seq, sequencing libraries of input and ChIP samples were prepared using Diagenode Microplex v3 (#C05010001) and UDIs (Set I: # C05010008; Set 2: #C05010009). Final libraries were quantified by qPCR using KAPA Illumina Library Quantification Kit (Roche #KK4854), with Agilent TapeStation used to estimate individual library sizes. Paired-end 50 bp or 100 bp sequencing was carried out by the Salk Next-Generation Sequencing Core on an Illumina NextSeq 2000 or NovaSeqX. The number of total and deduplicated sequencing reads is shown in **Table S2**.

*Antibodies:* Abflex anti-CTCF (Active Motif #91285; 4 µl per ChIP), anti-RAD21 (Abcam #ab992; 5 µl per ChIP), anti-STAG1 (Invitrogen #PA5-57115; 5 µl per ChIP), anti-STAG2 (Cell Signaling #5882; 5 µl per ChIP), anti-SCC-112/PDS5A (Bethyl #A300-089A; 10 µl per ChIP), anti-HA (Cell Signaling #3724 6 µl per ChIP), and anti-WAPL (Proteintech #16370-1-AP 5 µl per ChIP).

ChIP-seq analysis

*Alignment and processing:* Sequencing data from ChIP-seq was aligned using the Burrow-Wheeler Aligner (BWA)-MEM (*50*) to either the hg38 reference genome assembly alone or hg38 combined with mm10 (MM chromosome prefix) or dm6 (dm chromosome prefix) for experiments utilizing spike-in chromatin. Optical and PCR duplicates were removed using Picard MarkDuplicates (*51*) before samtools (*52*) was used to filter out reads with a MAPQ<30, improper pairs, and supplementary aligned reads. For sequencing data that included spike-in, samtools view -L was used to separate out reads that aligned to the primary chromosomes of hg38 from reads that aligned to the primary chromosomes of the spike-in species. Lastly, blacklisted regions (*53*) were then removed from hg38-aligned reads using BEDTools intersect (*54*).

*Peak calling:* Peaks from ChIP data were called using MACS2 (*55*, *56*), using input data as control. For CTCF ChIP-seq, peaks were called using a q-value cut-off of 0.001, whereas for RAD21 ChIP-seq, peaks were called with a q-value cut-off of 0.01. Aside from these modifications, peaks were called with the default settings, with file format set to BAMPE. For RAD21 ChIP-seq, we identified a set of high-confidence peaks based on peaks being called in ChIP-seq from the DMSO-treated conditions for both NIPBL-A2 and NIPBL-D7.

*Differential analysis:* High-confidence RAD21 peaks for the DMSO-treated condition (see above) were used to determine differential binding of RAD21 using DiffBind (*57*). Normalization was performed using the spike-in BAM files, with the DBA_DESEQ2 method and DBA_NORM_LIB. Differential peaks were determined using the DBA_DESEQ2 method.

*Meta-analysis and heatmaps:* deepTools bamCoverage was used to create BigWig files with sequencing data normalized using either a scale-factor (derived from DiffBind normFactor from spike-in chromatin) or RPKM (see figure legend), and using a bin size of 50 bp (*58*). computeMatrix was then used to capture signal values from the generated BigWig files using a given bed file (e.g. RAD21 ChIP-seq peaks).

*ChromHMM:* H3K9me3 (SRR15871779; input: SRR15871781) (*59*), H3K27me3 (SRR23280342; input: SRR18024435) (*60*), H3K36me3 (SRR23280340; input: SRR18024435) (*60*), H3K4me1 (SRR13259864; input: SRR13259866) (*61*), H3K4me3 (SRR13259862; input: SRR13259866) (*61*), H3K27ac (SRR10540139; input: SRR10540157) (*49*), and H3K9ac (SRR15871778; input: SRR15871781) (*59*) were downloaded from the Sequence Read Archive (SRA), with alignment and processing as described above. ChromHMM BinarizeBam was performed on the SRA data and our own CTCF ChIP-seq, using both the ChIP-seq and associated input files (*62*, *63*). We iteratively tested a range of models with ChromHMM LearnModel before settling on a 12-state model based on the fit of the ChIP data and interpretability of the output.

*Super-enhancer calling:* H3K27ac ChIP-seq in interphase hTERT RPE-1 cells was accessed from GEO (GSE141081) (*49*). ChIP and input sequencing data was aligned and processed as described above, utilizing the hg38/dm6 combined reference genome. Replicate ChIP and input bam files were merged before MACS2 callpeak was used to identify H3K27ac peaks, using a q-value cutoff of 0.001 (*55*, *56*). Rank Ordering of Super-Enhancers (ROSE) was performed using the called peaks, and the merged input and H3K27ac bam files (*64*, *65*). We tested multiple parameters (STITCHING_DISTANCE: 5000, 12500, 25000; TSS_EXCLUSION_ZONE_SIZE: 0, 2500, 5000), but focused our analyses on enhancers returned from a 12500 STITCHING_DISTANCE and 2500 TSS_EXCLUSION_ZONE_SIZE, which identified 1623 super-enhancers and 22,202 typical-enhancers.

*In situ* Hi-C

All Hi-C experiments were performed using the Arima Genomics Hi-C+ Kit, according to the manufacturer's instructions. Hi-C experiments were performed in independent clones presented which were used as experimental replicates (i.e. NIPBL-A2 and NIPBL-D7; RAD21-B1 and RAD21-B3), with the exception of the mitotic synchronization Hi-C experiments which were done in replicates using the NIPBL-D7 cell line. Briefly, cells were resuspended in DPBS and crosslinked with 2% methanol-stabilized formaldehyde (Fisher Scientific #F79500) for 10 minutes at room temperature. Stop Solution 1 was added to quench the formaldehyde, incubating first for 5 minutes at room temperature and then for 15 minutes on ice. Fixed cells were centrifuged, washed once with DPBS, and then flash frozen for storage at -80°C. Cells were lysed and nuclei permeabilized to allow for restriction digest of DNA before biotin fill-in ligation of proximal DNA ends. Samples were decrosslinked, digested with proteinase K, and DNA purified using Solid Phase Reverse Immobilization (SPRI) beads (Sera-Mag SpeedBead Carboxylate-Modified [E7] Magnetic Particles, Cytiva #45152105050250). Purified DNA was sonicated to ~400 bp using a Covaris M220 with 50 W power, 10% duty factor and 200 cycles per burst for 78 seconds, and then size-selected using SPRI beads. Based on quantification of DNA by Qubit, library preparation was either performed by KAPA HyperPrep (Roche #KK8502; for high input) based on Arima-provided instructions, or using the 2S Plus DNA Library Kit (IDT #10009878; for low input).

For KAPA HyperPrep, biotin labeled DNA was bound to Arima Enrichment Beads and used as input for end repair, dA-tailing, and ligation of TruSeq DNA Single Indexes (Illumina #20015960). Libraries were first quantified by qPCR using the KAPA Illumina Library Quantification Kit (Roche #KK4854) to estimate the required number of PCR cycles, before performing PCR amplification and SPRI bead clean-up. The final libraries were again quantified by qPCR using KAPA Illumina Library Quantification Kit (Roche #KK4854), and Agilent TapeStation used to estimate individual library sizes.

For 2S Plus DNA Library preparation, end repair and adaptor ligation were performed according to manufacturer's instructions. For biotin pull-down, 50 µl Dynabeads MyOne Streptavidin T1 beads (Invitrogen #65601) were washed 1X Tween Washing Buffer (5 mM Tris pH 8, 0.5 mM EDTA, 1 M NaCl, 0.05% Tween-20) and resuspended in 300 µl 2X binding buffer (10 mM Tris pH 8, 1 mM EDTA, 2 M NaCl) for addition to the end prepped/adaptor ligated DNA, with incubation for 15 minutes at room temperature. Beads were then washed twice at 55°C for two minutes using Tween Washing Buffer, washed once briefly using Low EDTA TE Buffer (from 2S Plus DNA Library Kit), and resuspended in Low EDTA TE Buffer. Unique combinatorial dual indexes (IDT #10009909) were added to each reaction before PCR amplification and SPRI bead clean-up. As above, final libraries were quantified by qPCR using KAPA Illumina Library Quantification Kit (Roche #KK4854), with Agilent TapeStation used to estimate individual library sizes.

To account for the accumulation of 4N cells with RAD21 depletion, we performed Hi-C mostly as described above. However, prior to decrosslinking, samples were incubated in propidium iodide staining buffer (10 µg/ml prodidium iodide (Abcam #ab14083), 100 µg/ml RNase (Thermo Scientific #EN0531), 2 mM $MgCl_2$, DPBS), and the G1/2N populations were sorted in bulk using BD Influx Cell Sorters in the Salk Flow Cytometry Core (see **Fig. S3A** for gating strategy used).

The sorted samples then reentered the Hi-C protocol starting with decrosslinking and proteinase K digestion.

Sequencing libraries first underwent a quality-control paired-end 37 bp run on Illumina MiniSeq, before paired-end 50 bp or 100 bp on Illumina NovaSeq6000, or paired-end 100 bp on NovaSeqX. Sequencing was carried out by the Salk Next-Generation Sequencing Core. The number of total and deduplicated sequencing reads, and cis/trans ratios are shown in **Table S3**.

Hi-C analysis

*Alignment and processing:* Hi-C data was aligned using a previously published in house pipeline (*66*). In brief, reads 1 and 2 were independently aligned using BWA-MEM to the hg38 reference genome assembly (*50*), before being combined post-alignment using "two_read_bam_combiner.pl". Optical and PCR duplicates were removed using Picard MarkDuplicates (*51*). A pairs format contact file was created using "bam_to_contacts_4DN_DCIC.pl". Contact files from experimental replicates were merged using "merge_contact_files.pl". A balanced 5 kb resolution cooler file was generated using "cooler cload" and "cooler balance", with "cooler zoomify" using the 5000N resolution subsequently used to create balanced multi-resolution cooler files (.mcool) (*67*). "Juicer pre" was used to create hic files from the contacts file, with "Juicer addNorm" subsequently used to add normalization from 5 kb (*68*).

*Chromatin loop calling and counts:* Juicer tools HiCCUPS (*68*) was used to call loops at the default resolutions and q-value cutoffs using Knight-Ruiz normalization. Loops called from the merged asynchronous, DMSO-treated NIPBL-A2/NIPBL-D7 Hi-C data was used as a canonical hTERT RPE-1 loop set. We used this loop set to access balanced loop counts from mcool files using "cooler dump" (*67*), expanding each loop bin in both directions along the horizontal and vertical to neighboring bins, and summing the balanced counts.

*Cluster analysis:* For cluster analysis, we first separated out cohesin-dependent and -independent chromatin loops based on a two-fold change in balanced loop count between RAD21-B1/B3 treated with DMSO- and dTAG$^{V}$-1. Balanced counts from asynchronous NIPBL-A2/D7 Hi-C for cohesin-dependent loops were then used as input into Cluster 3.0 (*69*, *70*) for K-means clustering based on Euclidean distance, using 100 repeats and six K-means clusters.

*Enhancer neighborhood:* Super-enhancers and typical-enhancers that were identified using ROSE (see above) that were within 1 Mb up- or downstream of a given TSS were classified as being within the "enhancer neighborhood". Enhancers that were within 10 kb of promoter-proximal regions were excluded. The contact frequency between promoter-proximal regions and neighborhood enhancers was visualized by pile-up analysis of observed/expected Hi-C data using cooltools (*71*), and by quantifying contact score using Chromosight with distance-dependence visualized by Loess-smoothening as described by Matthey-Doret, *et al.* (*72*) utilizing loess from scikit-misc.

*Additional analyses:* On-diagonal and off-diagonal pile-up analysis of observed/expected Hi-C data and contact vs. distance plots were performed using cooltools (*71*). For both of these, expected contact frequencies were determined from the input mcool file for each chromosome arm.

SLAM-seq

Cells were treated with 100 μM 4-thiouridine (4sU) for the amount of time mentioned (15 or 120 minutes), before being harvested in TRIzol (Invitrogen #15596018). RNA was prepared under red light using the Direct-zol RNA Miniprep Kit (Zymo #R2052) and then treated with iodoacetamide using the SLAMseq Kinetics Kit – Anabolic Kinetics Module (Lexogen #061), according to manufacturer's instructions. For the mitosis synchronization experiments, the RNA was then purified by ethanol precipitation, while for asynchronous depletion experiments, we used the RNA Clean & Concentrator-5 kit (Zymo #R1013). RNA integrity was determined using Agilent TapeStation. To maximize the recovery of nascent RNA, library preparation was performed using Zymo-Seq RiboFree Total RNA Library Kit (Zymo #R3003), according to manufacturer's instructions. Library quantification was performed by qPCR using KAPA Illumina Library Quantification Kit (Roche #KK4854), with library sizes estimated using Agilent TapeStation.

Next-generation sequencing was performed by the Salk Next-Generation Sequencing Core on an Illumina NovaSeq6000 using paired-end 50 bp or 100 bp. The number of total and deduplicated sequencing reads is shown in **Table S4**.

SLAM-seq analysis

*Alignment and processing:* The previously described SLAM-DUNK pipeline (*73*) for alignment of SLAM-seq sequencing data and quantification of T→C conversions in nascent RNA utilizes the alignment tool NextGenMap. Although this is a reasonable alignment approach when performing the 3' mRNA sequencing (Quant-seq) that has previously been employed for SLAM-seq library preparation (*32*), NextGenMap is unable to perform the gapped alignments that are a preferred for RNA sequencing reads and is therefore incompatible with the Zymo-Seq RiboFree Total RNA Library Kit (Zymo #R3003) utilized here. Instead, we implemented our own analysis pipeline built around HISAT-3N (*74*), a three-nucleotide alignment strategy more suitable for SLAM-seq data derived from total RNA sequencing.

First, to account for potential single nucleotide polymorphisms that may be present in the sequencing data and have the potential to cause false positives in the identification of nascent RNA, we replaced homozygous and heterozygous SNPs (*75*) with N in the hg38 reference genome. This modified reference genome was used to build a HISAT-3N index with the T→C base change appropriate for SLAM-seq. Directional mapping was subsequently performed using HISAT-3N. For the Zymo-Seq RiboFree Total RNA Library Kit (Zymo #R3003) utilized here, read 1 is anti-sense and read 2 is sense. We found that, for T→C conversions to be appropriately identified in paired-end sequencing from these libraries, the reads had to be switched for alignment with HISAT-3N (i.e. -1 read2.fastq.gz -2 read1.fastq.gz) while using '--rna-strandness FR'. As recommended by the Zymo-Seq RiboFree Total RNA Library Kit protocol, we also trimmed 10 bp from the 5' end of reads during the alignment. HISAT-3N returns aligned reads that carry Yf, Zf, and YZ SAM tags that mark the number of T→C conversions in the read, the number of unconverted Ts in the read, and whether the read mapped to the positive or negative strand, respectively. Following alignment, optical and PCR duplicates were removed using Picard MarkDuplicates (*51*).

*Gene quantification:* To quantify nascent and total transcripts for a given gene, we created a custom Python script utilizing HTSeq (*76*). In this, we filter out unmapped, secondary, and supplementary reads, and those with a MAPQ<10 With remaining reads that unambiguously align to a gene from Gencode v25, those whereby the Yf field is greater than 0, indicating at least one T→C conversion, are counted as nascent, while all unambiguously aligning reads, regardless of value in the Yf field, are counted as total.

*Differential gene expression analysis:* Differentially expressed genes were identified using DESEQ2 (*77*). To account for different levels of nascent transcripts between samples, DESEQ2 "sizeFactors" derived from total RNA read counts were used to normalize nascent (T→C) read counts. A low expression filter was then applied to remove lowly expressed genes from both the nascent and total gene sets; genes whereby the minimum number of reads was exceeded in the minimum number of replicates in a given comparison were kept (e.g. 3 replicates per condition: 3 samples in the comparison must exceed the minimum), while genes not meeting this criteria were excluded (minimum reads for inclusion; total RNA: 20 reads; nascent 15min 4sU: 2 reads, nascent 120min 4sU: 20 reads). Tables containing FPKM information for plotting was outputted from DESEQ2 prior to filtering.

*Nascent read BigWig tracks:* BamTools (*78*) was used to identify reads containing at least one T→C conversion (Yf:>0), and based on this we extracted read pairs whereby at least one read contained a T→C conversion using SAMtools (*52*). We then used the reciprocal of the sizeFactors for total RNA from DESEQ2 to normalize nascent read pairs in a BigWig format using deepTools bamCoverage (*58*).

*Gene ontology (GO) analysis and gene set enrichment analysis (GSEA):* TopGO was used to perform gene ontology analysis for biological process using the "elim" algorithm (*79*). Note that p-value adjustment is not recommended for this algorithm. Pre-ranked GSEA against the MSigDB hallmark gene set (*80*) was performed on a $\log_2$FC-sorted list of genes returned by DESEQ2 using the command line version of the UCSD/Broad GSEA tool (*81*).

Statistics

With the exception of in-built statistical testing for DESEQ2, GO analysis, and GSEA, statistical tests were carried out using SciPy in Python (*82*). See figure legend for the specific statistical test used.

Additional tools

Plotting of Hi-C heatmaps was performed using cooler (*67*), and BigWig and Virtual4C tracks using CoolBox (*83*). Figures were largely generated using Matplotlib (*84*) and seaborn (*85*). Flow cytometry gating during collection was performed using FACSDiva (*86*), while gating and quantification of flow cytometry data for visualization was performed using FlowJo (*87*).

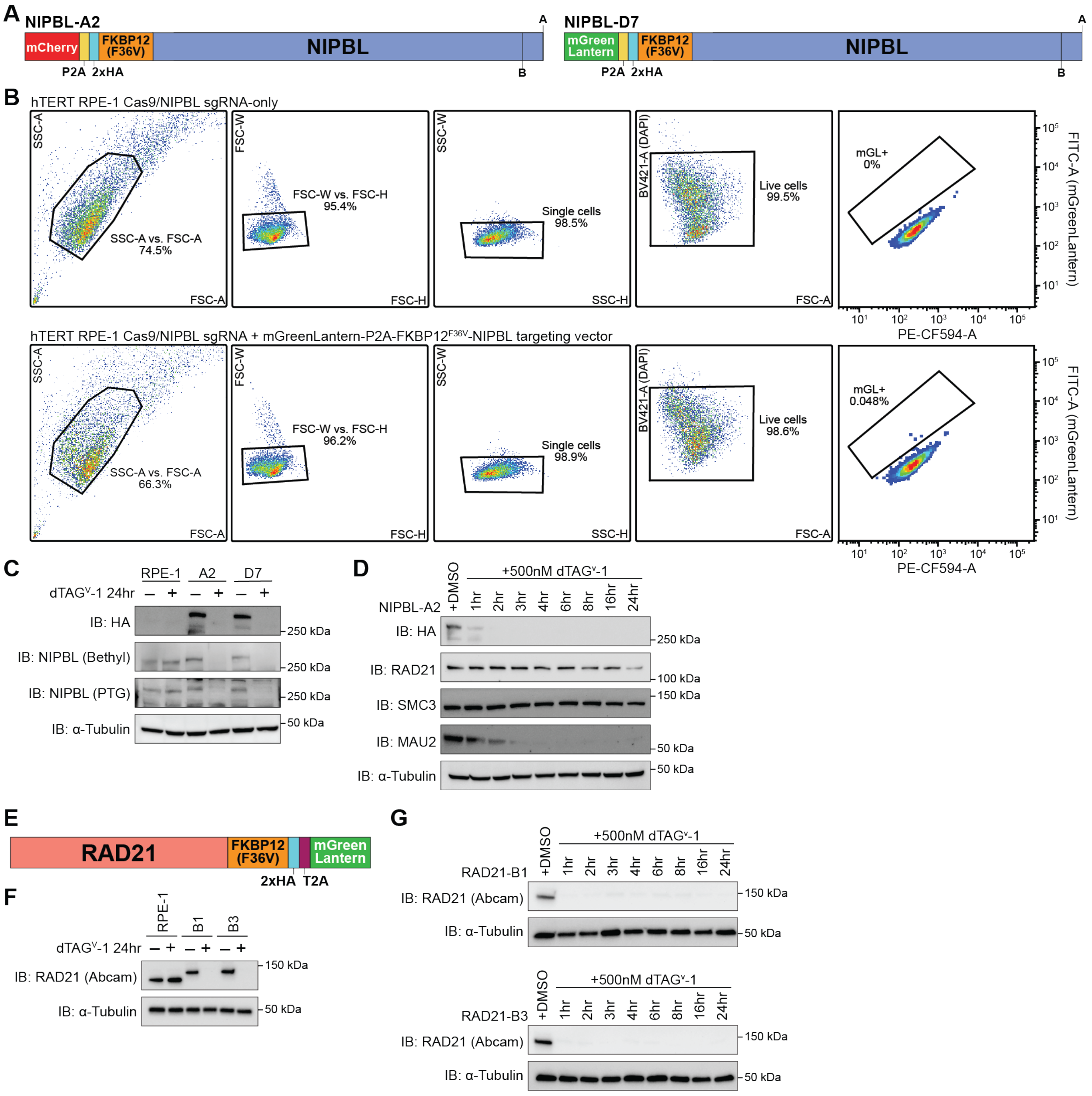


**Fig. S1 (related to Figure 1): Acute depletion of NIPBL and RAD21 from hTERT RPE-1 cells.** (**A**) Schematics of N-terminal FKBP12$^{F36V}$-NIPBL fusions engineered into hTERT RPE-1 cells using CRISPR/Cas9-directed homologous recombination. N-terminal tagging was employed to maximize degradation of NIPBL isoforms (A and B). The NIPBL-A2 and NIPBL-D7 clones are homozygous for a P2A-linked mCherry and P2A-linked mGreenLantern, respectively. (**B**) Example gating strategy used for fluorescence activated cell sorting of hTERT RPE-1 cells transfected with Cas9/*NIPBL* sgRNA, with or without a targeting vector containing mGreenLantern-P2A-FKBP12$^{F36V}$ flanked by *NIPBL* homology arms. Cells were separated from debris by gating in SSC-A vs. FSC-A, then single cells were obtained by sequential gating on FSC-W vs. FSC-H and SSC-W vs. SSC-H. Lastly, live cells were gated based on absence of DAPI, before mGreenLantern (mGL)-positive cells were sorted based on absence in sgRNA-only and presence in sgRNA + targeting vector. (**C**) Whole cell extracts were harvested from parental hTERT RPE-1 and the engineered NIPBL-A2 and NIPBL-D7 cell lines treated with DMSO or dTAG$^{V}$-1 for 24 hours. The shift in molecular weight of NIPBL in the engineered cell lines is demonstrated by western blot using two different commercially available polyclonal antibodies raised against NIPBL (PTG: 18792-1-AP; Bethyl: A301-779A). (**D**) Western blot of whole-cell lysates from NIPBL-A2 cells treated with DMSO for 24 hours or dTAG$^{V}$-1 for varying lengths of time to determine rate of HA-NIPBL depletion. α-Tubulin is shown as a loading control. (**E**) Schematic of C-terminal RAD21-FKBP12$^{F36V}$ fusion engineered in hTERT RPE-1 cells using CRISPR/Cas9-directed homologous recombination. The RAD21-B1 and RAD21-B3 clonal cell lines are homozygous for T2A-linked mGreenLantern. (**F**) Western blot of parental hTERT RPE-1, RAD21-B1, and RAD21-B3 cells treated with DMSO or dTAG$^{V}$-1 for 24 hours. (**G**) Lysates for western blot were harvested from RAD21-B1 (top) and RAD21-B3 (bottom) cells treated with DMSO for 24 hours or dTAG$^{V}$-1 for increasing lengths of time.

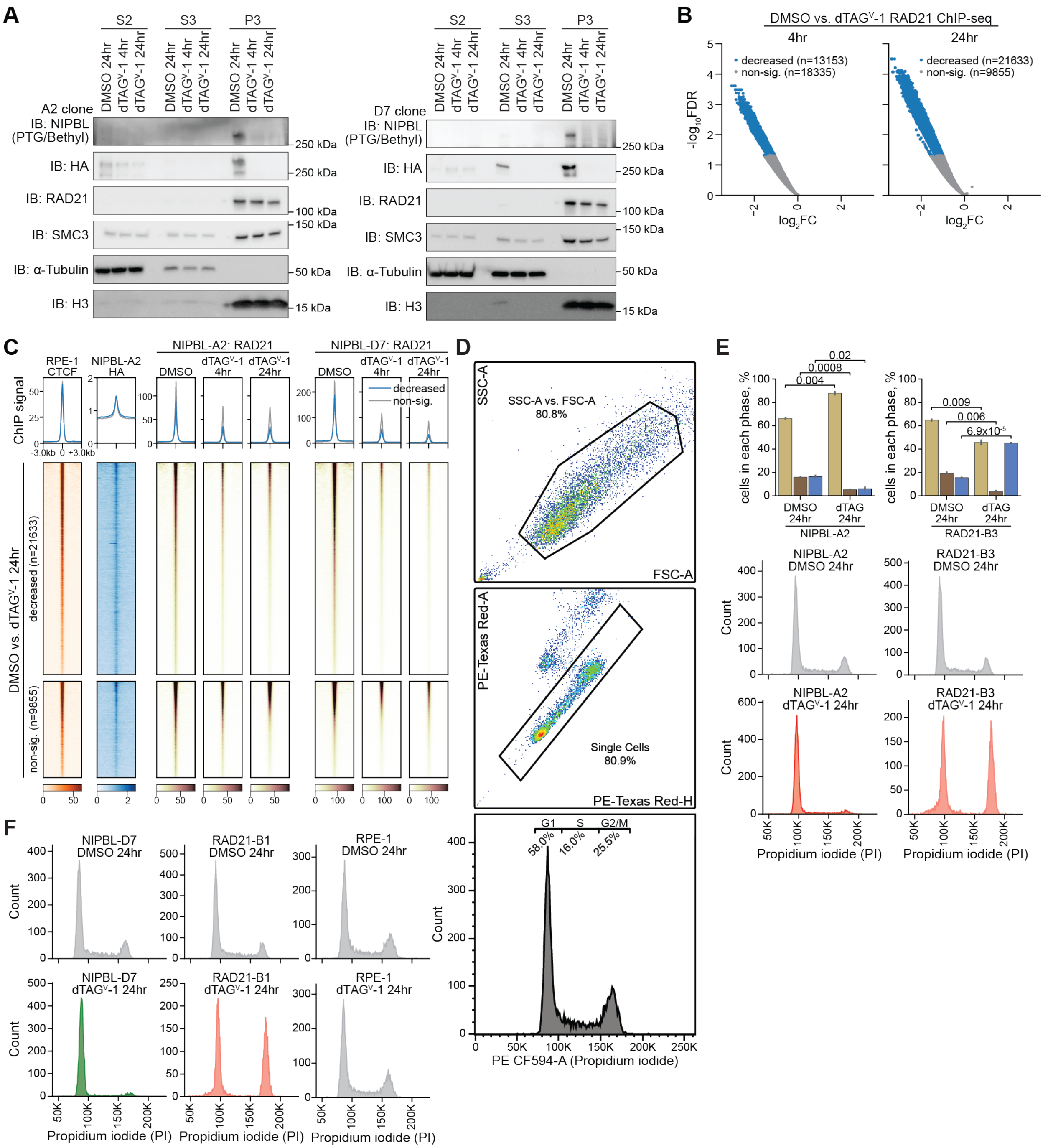


**Fig. S2 (related to Figure 1): NIPBL is necessary for most, but not all RAD21 ChIP-seq peaks.** (**A**) Chromatin fractionation was used to separate out cytoplasmic (S2), nucleoplasmic (S3), and chromatin (P3) fractions from DMSO- or dTAG$^{V}$-1-treated NIPBL-A2 (left) and NIPBL-D7 (right) cells. Two different commercially available polyclonal antibodies raised against NIPBL (PTG: 18792-1-AP; Bethyl: A301-779A) were combined to improve detection of NIPBL in western blotting. Fractionation success was determined by the absence of α-Tubulin and presence of H3 in the P3 fraction, and vice versa in the S2 and S3 fractions. (**B**) Volcano plots from differential analysis of high-confidence RAD21 ChIP-seq peaks with NIPBL depletion for 4 hours (left) and 24 hours (right). RAD21 ChIP-seq from NIPBL-A2 and NIPBL-D7 were treated as biological replicates. No significantly increased peaks were observed, but significantly decreased peaks (FDR < 0.05) are highlighted in blue. (**C**) Based on differential analysis of RAD21 ChIP-seq in FKBP12$^{F36V}$-NIPBL cells with a 24-hour dTAG$^{V}$-1 treatment, we show scale factor-normalized RAD21 ChIP signal at significantly decreased and non-significantly changed peaks. RPKM-normalized ChIP signal for HA-NIPBL and CTCF is also shown. RAD21 ChIP-seq from NIPBL-A2 and NIPBL-D7 cell lines were treated as replicates for differential analysis. (**D**) Example flow cytometry gating strategy used for determining cell cycle distribution by propidium iodide staining. Cells were separated from debris by gating in SSC-A vs. FSC-A, then single cells were selected based PE-Texas Red-A vs. PE-Texas Red-H. Depending on the flow cytometer used, propidium iodide was detected using either the PE-Texas Red or PE-CF594 channel. (**E**) Quantification (top) and examples (bottom) of cell cycle distributions by propidium iodide staining of NIPBL-A2 and RAD21-B3 cells treated with DMSO or dTAG$^{V}$-1, as measured by flow cytometry. Independent, two-sided t-tests were performed on cell cycle distributions for each cell line using two replicates. (**F**) Example cell cycle distributions from propidium iodide staining of NIPBL-D7, RAD21-B1, and parental RPE-1 cells treated with DMSO or dTAG$^{V}$-1 for 24 hours, as measured by flow cytometry. Quantification is shown in Figure 1C.

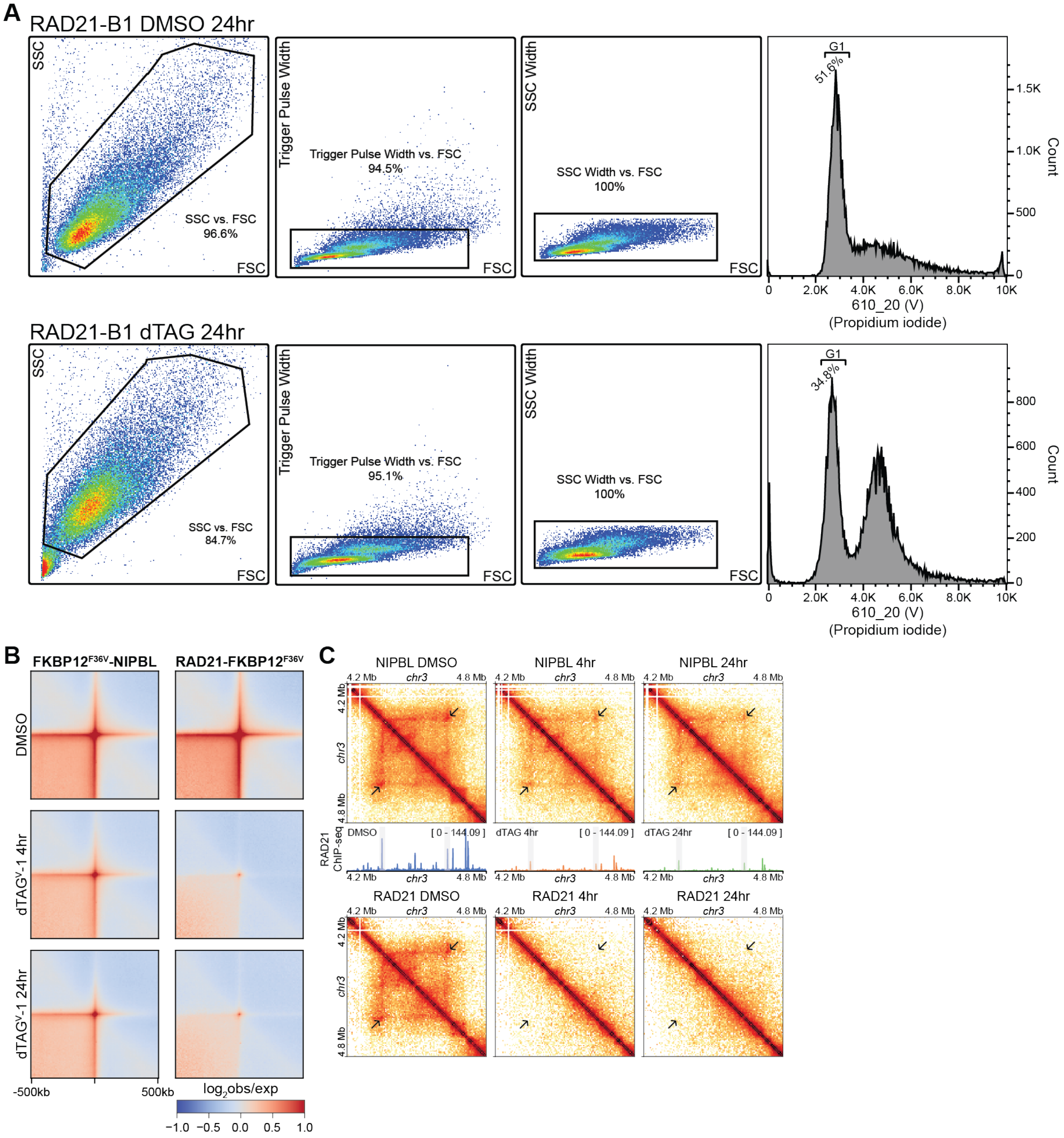


**Fig. S3 (related to Figure 1): Consequences of NIPBL and RAD21 depletion on genome organization and nascent transcription.** (**A**) Gating strategy used for fluorescence activated cell sorting of 2N RAD21-B1/B3 cells for Hi-C. Cells first underwent initial steps for Hi-C, including crosslinking, restriction digestion, and proximity ligation, before being stained with propidium iodide for sorting. Post-sorting, cells were decrosslinked and sequencing libraries prepared. (**B**) Off-diagonal pile-up analysis showing log-transformed observed/expected contact frequency from Hi-C at chromatin loops called from control, DMSO-treated FKBP12$^{F36V}$-NIPBL cells. (**C**) Example heatmaps showing the consequences of NIPBL (top) and RAD21 (bottom) depletion for 4 and 24 hours at the same locus. Beneath the NIPBL heatmaps are scale factor-normalized RAD21 ChIP-seq tracks for the corresponding treatment in NIPBL-D7 cells. A chromatin loop of interest is highlighted with an arrow in the heatmaps, while the loop anchors in the ChIP-seq tracks are marked a grey box.

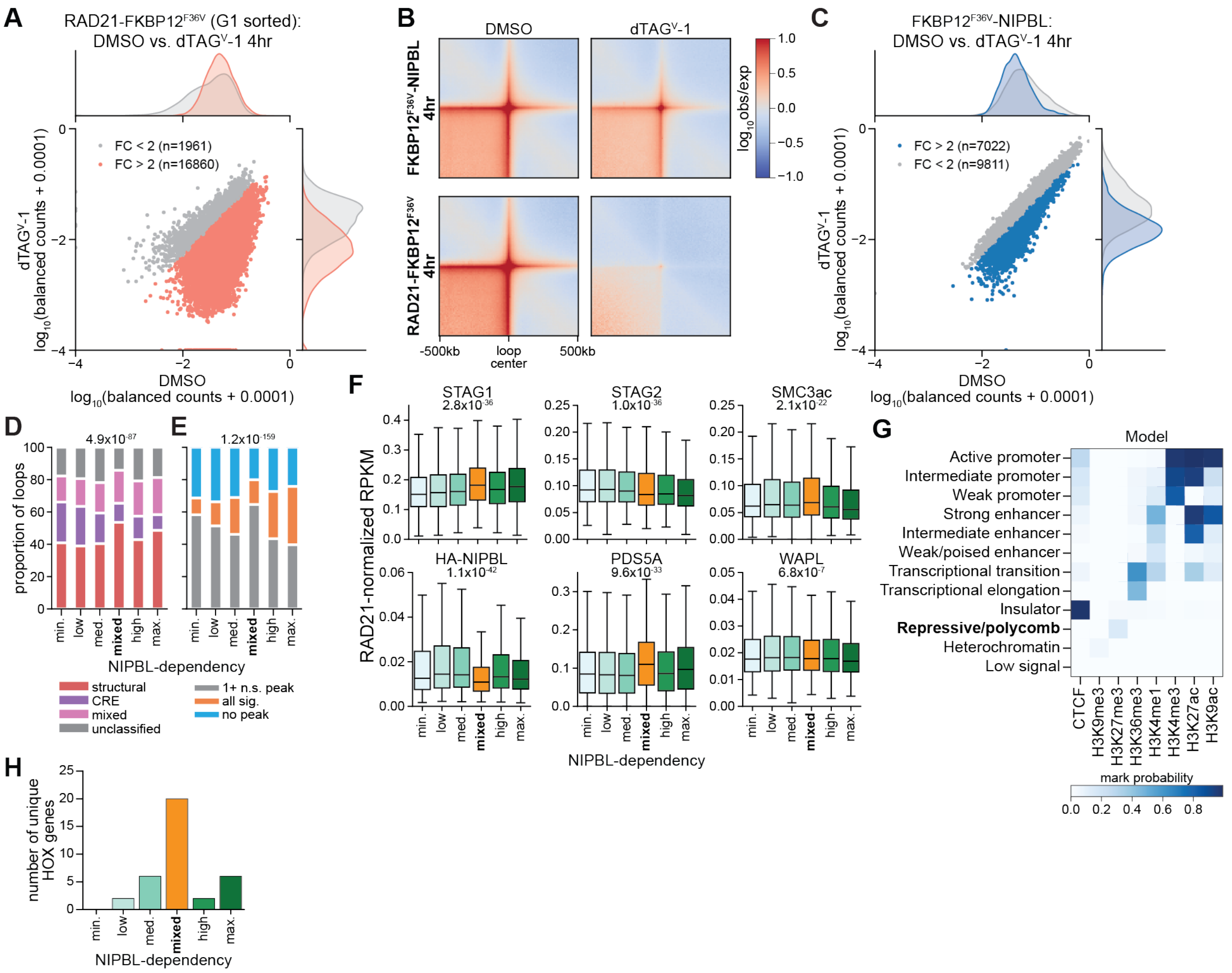


**Fig. S4. (related to Figure 2): Chromatin loops show differential dependence on NIPBL and RAD21.** (**A**) Scatter plot showing the log-transformed balanced counts between DMSO and dTAG$^{V}$-1-treated conditions with RAD21 depletion. Chromatin loops that showed a fold-change of at least two are highlighted. A kernel density estimate (KDE) plot is shown on each axis to represent the distribution of the datapoints on that axis. To each balanced count value, 0.0001 was added prior to log-transformation to allow for plotting of zero values. (**B**) Observed/expected pile-up analysis at a 10 kb resolution at RAD21-dependent chromatin loops (RAD21 FC > 2) from Hi-C performed on DMSO- and dTAG$^{V}$-1-treated FKBP12$^{F36V}$-NIPBL and RAD21-FKBP12$^{F36V}$ cells. (**C**) Scatter plot of RAD21-dependent chromatin loops (RAD21 FC > 2) showing the log-transformed balanced counts between DMSO and dTAG$^{V}$-1-treated FKBP12$^{F36V}$-NIPBL cells. Chromatin loops that showed a fold-change of at least two are highlighted. A kernel density estimate (KDE) plot is shown on each axis to represent the distribution of the datapoints on that axis. To each balanced count value, 0.0001 was added prior to log-transformation to allow for plotting of zero values. (**D**) Classification of chromatin loops based on their association with RAD21/CTCF ChIP-seq peaks (structural), cis regulatory element (CRE), or a mix of both. P-value is from a chi-squared test. (**E**) Relationship between clusters and differential RAD21 ChIP-seq peaks with depletion of NIPBL for 4-hours. (**F**) Boxplot showing the RPKM at RAD21 ChIP-seq peaks that intersect the chromatin loop anchors from ChIP-seq against cohesin subunits and accessory proteins. For each chromatin loop, the mean RPKM was taken from all RAD21 ChIP-seq peaks across both anchors, and was normalized to the mean RPKM of RAD21 at the same anchors to account for differences in RAD21 binding at different loops. Differences in chromatin binding at the clusters was tested using a Kruskal-Wallis test. (**G**) Model of chromatin states in hTERT RPE-1 cells generated using ChromHMM. Mark probability is from the ChromHMM emission parameters and estimates the likelihood that a given mark would be found in that state. (**H**) Number of unique HOX genes with promoter-proximal regions (TSS -1 kb/+0.5 kb) intersecting chromatin loop anchors from the different clusters. Each cluster of chromatin loops are associated with different HOX clusters, specifically low-dep. overlaps HOXB, med.-dep. overlaps HOXA, HOXC, and HOXD, mixed-dep. overlaps HOXA, HOXB, HOXC, and HOXD, high-dep. overlaps HOXD, and max.-dep. overlaps HOXA, HOXB, and HOXC.

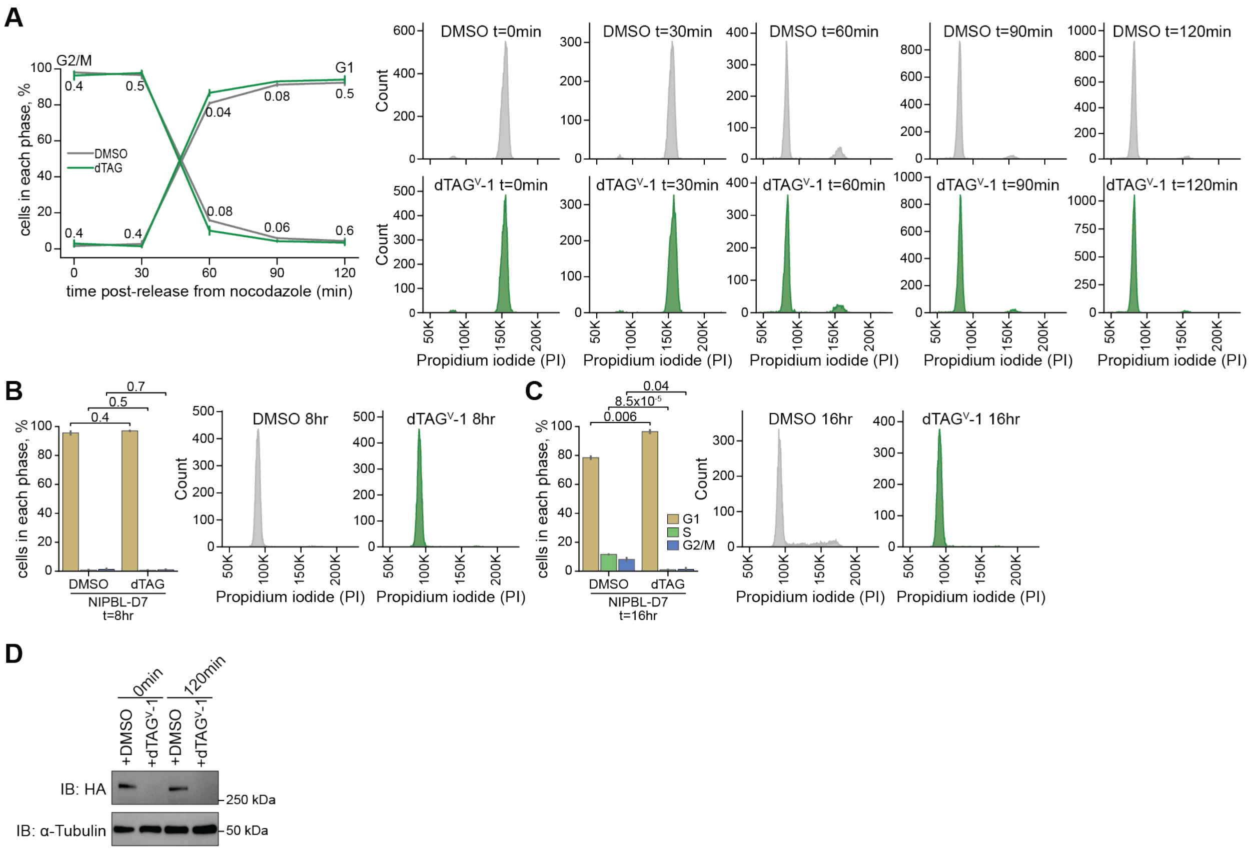


**Fig. S5. (related to Figure 3): Depletion of NIPBL during mitotic exit. (A)** Quantification (left) and examples (right) of cell cycle distributions by propidium iodide staining of NIPBL-D7 cells treated with DMSO or dTAG$^{V}$-1 during nocodazole synchronization and mitotic exit, as measured by flow cytometry. Cells were collected at t=0min following a 2-hour nocodazole synchronization with DMSO/dTAG$^{V}$-1 treatment, and at 30-minute intervals following release. Independent, two-sided t-tests were performed between the DMSO and dTAG$^{V}$-1-treated conditions using two replicates. Note that at t=60min we detect a comparatively high proportion of "doublets" under both treatment conditions, likely reflecting cells undergoing cytokinesis. (**B & C**) Quantification (left) and examples (right) of cell cycle distributions by propidium iodide staining of NIPBL-D7 cells treated with DMSO or dTAG$^{V}$-1 during nocodazole synchronization and mitotic exit, as measured by flow cytometry. Cells were harvested 8 hours (**B**) and 16 hours (**C**) post-release from nocodazole. Independent, two-sided t-tests were performed between the DMSO and dTAG $^{V}$-1-treated conditions using two replicates. (**D**) Western blot of cell lysates harvested at t=0min and t=120min following nocodazole synchronization and release with DMSO/dTAG$^{V}$-1 treatment of the NIPBL-D7 cell line.

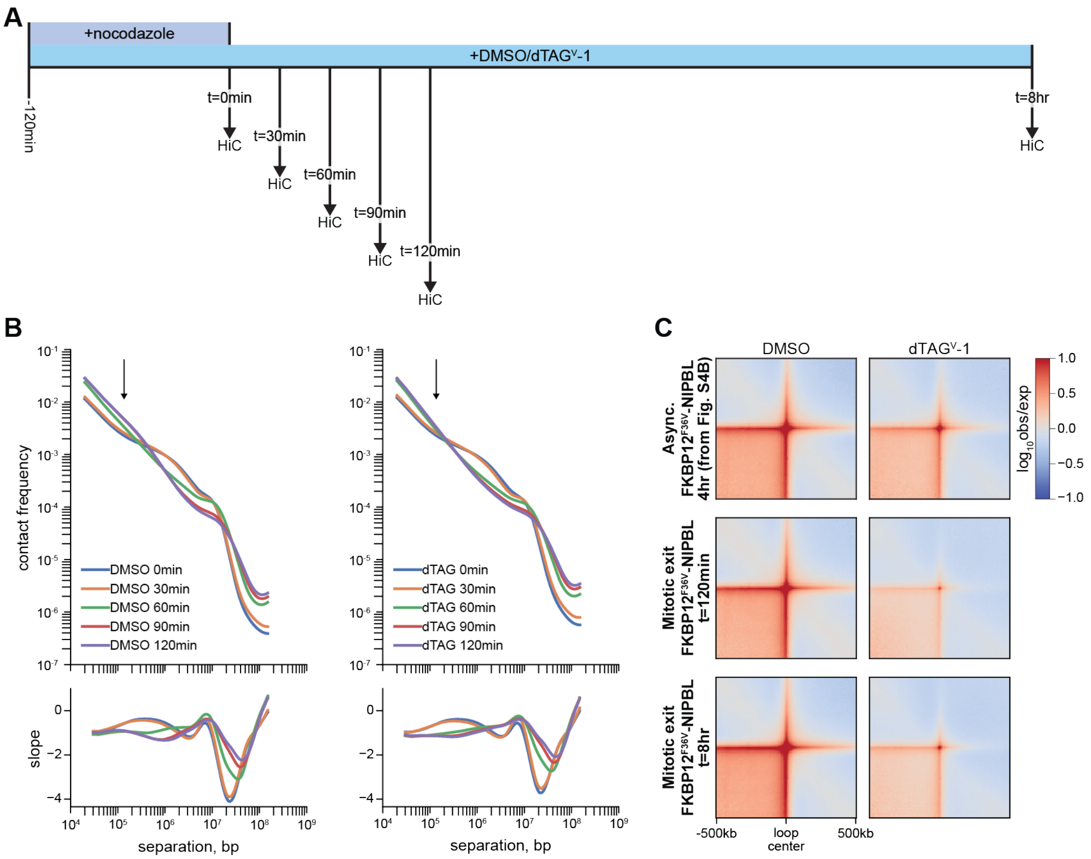


**Fig. S6. (related to Figure 3): NIPBL is more critical to the post-mitotic establishment of chromatin loops than their maintenance in interphase cells.** (**A**) Schematic outlining the approach to perform Hi-C with depletion of NIPBL during mitotic exit. Cells were synchronized to prometaphase using a double thymidine block (14 hours thymidine – 10 hour release – 14 hour thymidine – 8 hour release) and nocodazole treatment. DMSO/dTAG$^{V}$-1 was added simultaneous with nocodazole. (**B**) Genome-wide contact frequency over genomic distance at a 10 kb resolution from Hi-C for DMSO- and dTAG$^{V}$-1 NIPBL-D7 cells during mitotic exit. Cells were collected at 30-minute intervals starting with nocodazole-arrested cells (t=0min). Data from replicate experiments were combined for this analysis. The arrows highlight the region of these plots that reflect chromatin loops. (**C**) Observed/expected pile-up analysis of Hi-C data at a 10 kb resolution of cohesin-dependent chromatin loops identified in Figure S4A.

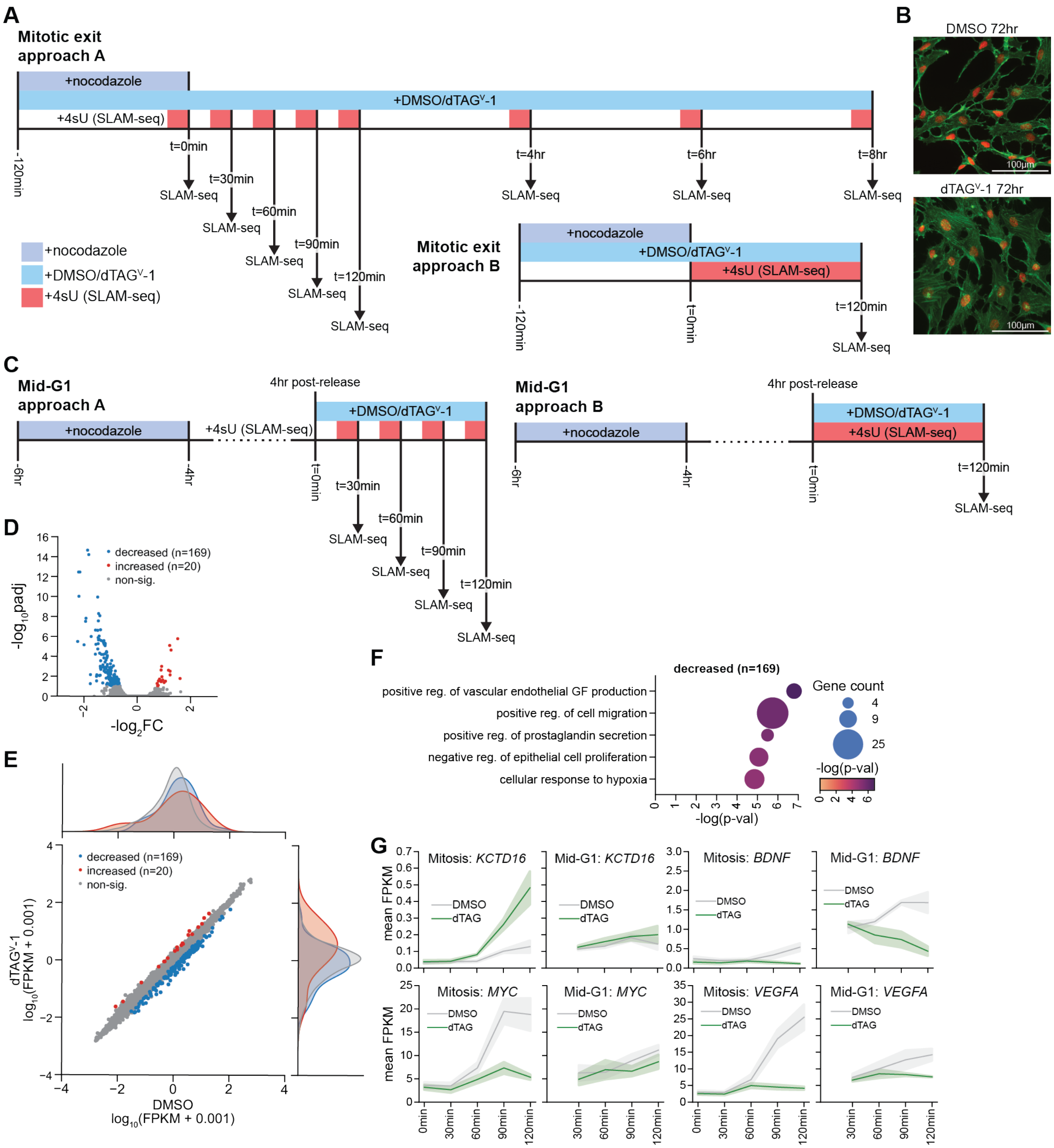

**Fig. S7. (related to Figure 4): Approaches used to interrogate the contribution of NIPBL to gene regulation during mitotic exit and G1.** (**A**) Schematic outlining the two approaches used to investigate the consequences of NIPBL depletion during mitotic exit. For both approaches, cells first underwent a double thymidine block (14 hours thymidine – 10 hour release – 14 hour thymidine – 8 hour release) prior to nocodazole treatment. These approaches differ in the length of 4sU treatment for SLAM-seq, with approach A utilizing short 15 minute labeling periods, whereas approach B utilized a 120min long labeling period corresponding to the extent of the release period. (**B**) Morphology of NIPBL-A2 cells treated with DMSO or dTAG$^{V}$-1 for 72 hours. Cells were stained with phalloidin (actin; green) and DAPI (DNA; red). (**C**) Schematic outlining the two approaches used to investigate the consequences of NIPBL depletion during G1. As described above, cells were first synchronized with double thymidine block (14 hours thymidine – 10 hour release – 14 hour thymidine – 8 hour release) before a 2 hour nocodazole treatment. Cells were then released for 4 hours before addition of DMSO or dTAG$^{V}$-1 for 2 hours. For SLAM-seq, cells were treated with 4sU for either 15 minutes prior to harvesting (approach A) or for 2 hours over the length of the DMSO/ dTAG$^{V}$-1 treatment (approach B). (**D**) Volcano plot from differential expression analysis of nascent transcripts between DMSO- and dTAG$^{V}$-1-treated NIPBL-D7 cells based on a 120min 4sU labeling period beginning four hours post-release from nocodazole (mid-G1 approach B, Figure S7C). Significantly changed ($|log_2FC|>0.585$ and $padj<0.1$) nascent transcripts are highlighted. Differential analysis was performed using three independent replicates. (**E**) Scatter plot showing the log-transformed FPKM values for nascent transcripts in the DMSO- and dTAG$^{V}$-1-treated NIPBL-D7 cells following a 120min 4sU labeling period in the middle of the G1 cell cycle phase. Significantly changed transcripts ($|log_2FC|>0.585$ and $padj<0.1$) are highlighted. A kernel density estimate (KDE) plot is shown on each axis to represent the distribution of the datapoints on that axis. To each FPKM value, 0.001 was added prior to log-transformation to allow for plotting of zero FPKM values. (**F**) Biological process gene ontology analysis was performed on the significantly decreased nascent transcripts between DMSO and dTAG$^{V}$-1 following a 120min 4sU labeling period in the middle of the G1 cell cycle phase. The top 5 categories are shown for each analysis, with the P-values derived using the 'elim' algorithm from TopGO. (**G**) As in Figure 4J, but showing line plots for additional example differentially expressed genes, plotting the mean FPKM and standard error based on mitotic exit and mid-G1 approach A. All shown genes were significantly changed during mitotic exit, but only *BDNF* and *VEGFA* were also significantly changed in mid-G1. For "mitosis", the x-axis is time post-release from nocodazole, while for "mid-G1", the x-axis is time post-addition of DMSO/ dTAG$^{V}$-1.

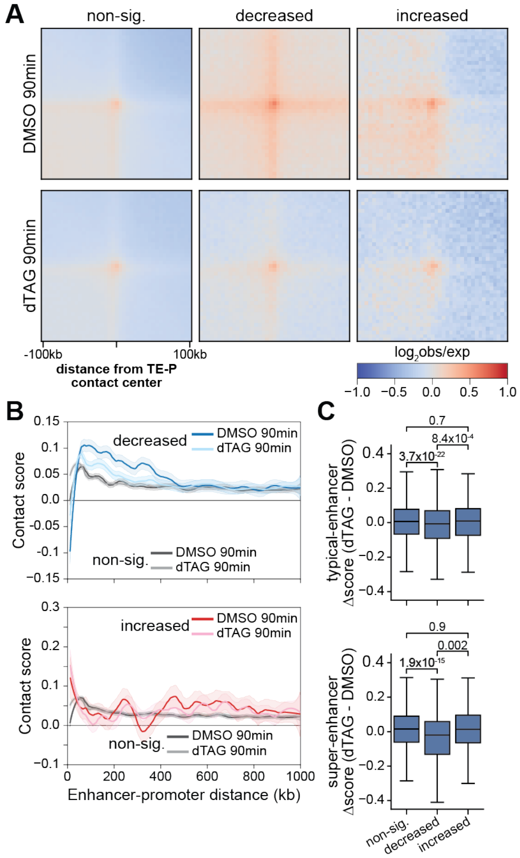


**Fig. S8. (related to Figure 5): Typical-enhancer neighborhood and sensitivity to NIPBL depletion.** (**A**) As in Figure 5B, but for the typical-enhancer neighborhood. (**B**) Loess-smoothened relationship with 95% confidence interval at a 5 kb resolution between distance and contact frequency between typical-enhancers and promoter-proximal regions (TSS -1 kb/+500 bp). For plotting of non-significant contact scores, 40000 enhancer-promoter pairs were randomly selected for each of the plots. (**C**) Quantification of the difference in contact scores for typical-enhancers (top) and super-enhancers (bottom) that are within 500 kb of a TSS from Figure 5C and Figure S8B. P-values were determined using a two-sided Wilcoxon rank-sum test.

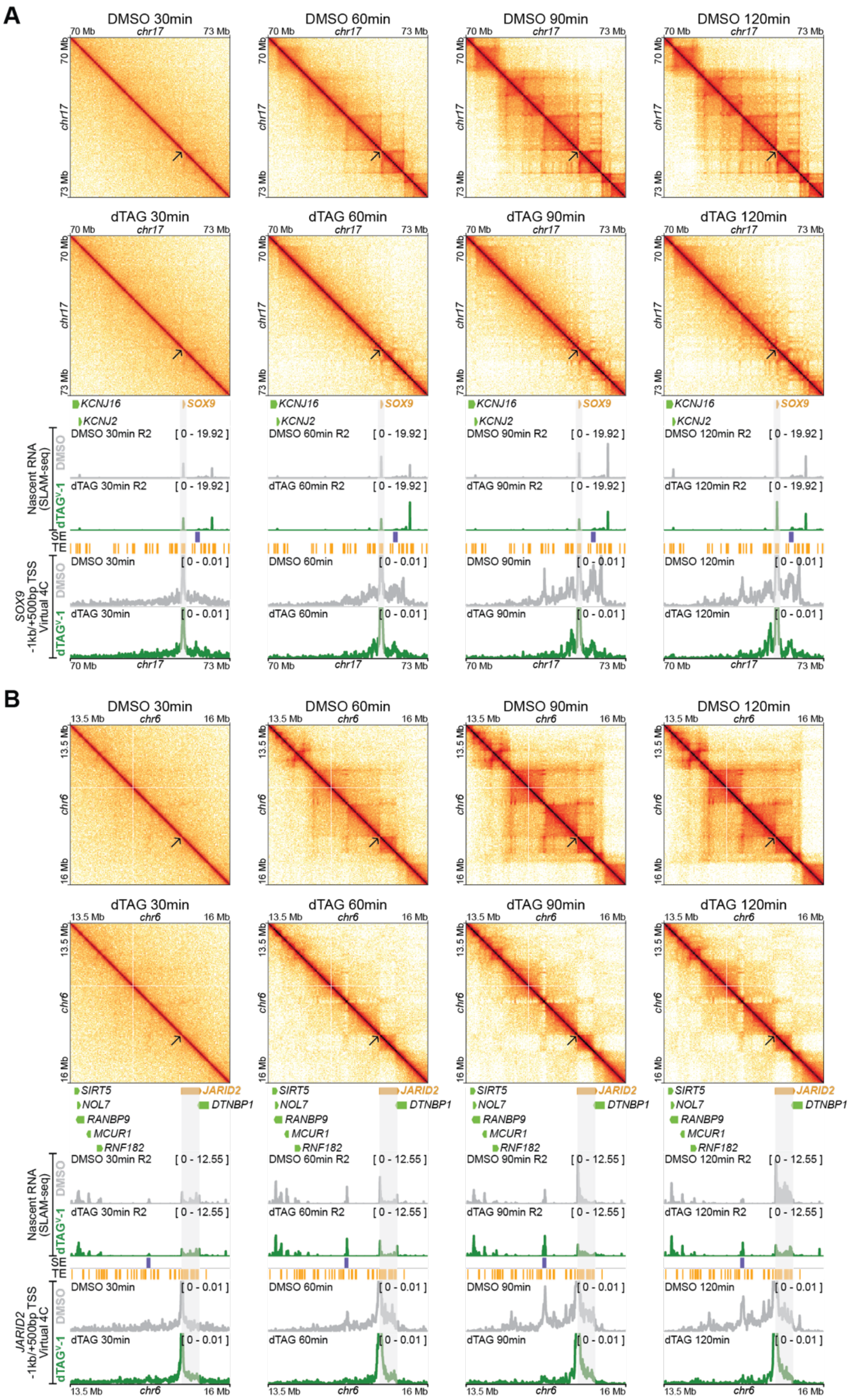


**Fig. S9. (related to Figure 5): Examples of genes sensitive to NIPBL depletion during mitotic exit.** *SOX9* (**A**) and *JARID2* (**B**) are examples of genes that are significantly decreased with NIPBL depletion during mitotic exit. Shown are Hi-C heatmaps (top: DMSO; bottom: dTAG), sequencingdepth normalized BigWig tracks of read pairs whereby at least one read contains a T→C conversion (i.e. nascent reads), and Virtual 4C anchored at promoter-proximal regions (TSS -1 kb/+500 bp) for both DMSO and dTAG-treated samples at t=30min, 60min, 90min, and 120min. The marked TE and SE sites are the same across all plots. The arrow on the Hi-C heatmaps marks the transcription start site (TSS), and the grey bar on the tracks below marks the gene location.

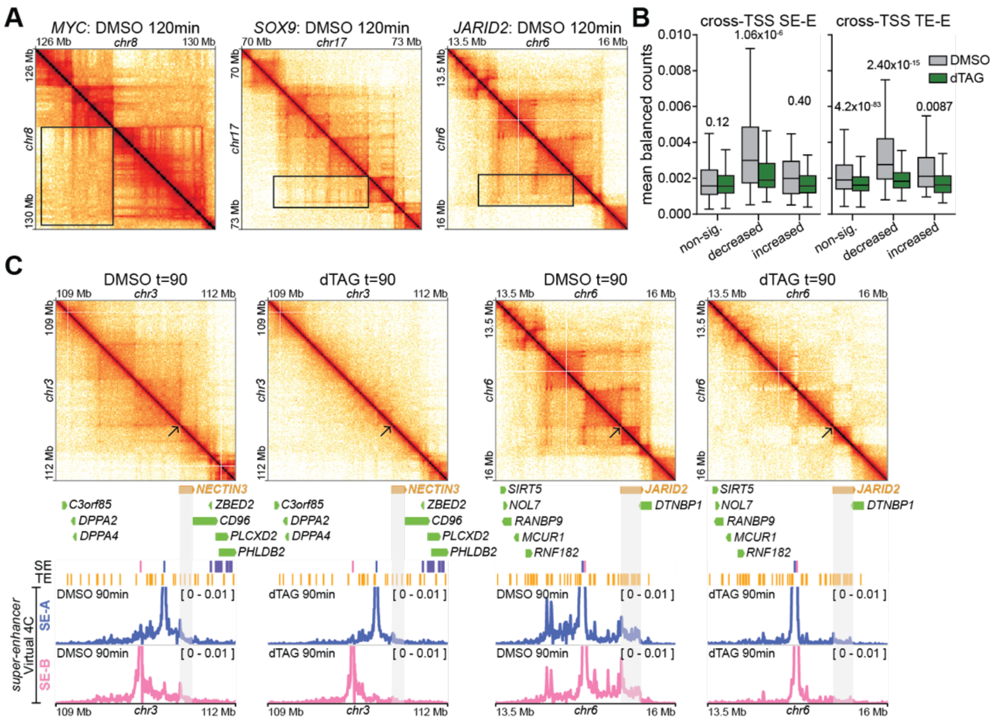


**Fig. S10. (related to Figure 5): Spatial proximity of pairs of neighborhood enhancers.** (**A**) Cross-TSS connectivity at example genes sensitive to NIPBL depletion during mitotic exit. The top-right corner of each box corresponds to the TSS of each gene. (**B**) Balanced counts between super-enhancers (left) or typical-enhancers (right) and enhancers on the opposite side of a TSS. Each enhancer that was between 15 kb and 1 Mb from a TSS was assigned to three 5 kb bins, centered on the enhancer before determining the contact frequency based on a balanced matrix. For each enhancer-enhancer pair, balanced counts from the different bin pairs were summed, and for each gene, the mean of the different, non-zero enhancer-enhancer pairs was taken before plotting. P-values were determined using a two-sided Wilcoxon rank-sum test. (**C**) Examples of TSS insulation at a non-significantly changed gene (NECTIN3) and reduced TSS insulation at a significantly decreased gene (JARID2). For each gene we show Virtual 4C anchored at two different super-enhancers (SE-A: blue, SE-B: pink) that are upstream of the TSS. The arrow marks the transcription start site (TSS).

**Table S1.**

Cloning primers used introduce sgRNAs against NIPBL and RAD21 into PX330, and sequencing primers used to validate insertion of targeting vector payload into the NIPBL and RAD21 loci.

| Name | Sequence | Description |
|---|---|---|
| **NIPBL-sgRNA_F** | caccgAGAGTAGTAATGGGGACATG | Forward primer used to introduce sgRNA (AGAGTAGTAATGGGGACATG) targeting NIPBL into PX330. |
| **NIPBL-sgRNA_R** | aaacCATGTCCCCATTACTACTCTc | Reverse primer used to introduce sgRNA (AGAGTAGTAATGGGGACATG) targeting NIPBL into PX330. |
| **RAD21-sgRNA_F** | caccgCCTTATATAATATGGAACCT | Forward primer used to introduce sgRNA (CCTTATATAATATGGAACCT) targeting RAD21 into PX330. |
| **RAD21-sgRNA_R** | aaacAGGTTCCATATTATATAAGGc | Reverse primer used to introduce sgRNA (CCTTATATAATATGGAACCT) targeting RAD21 into PX330. |
| **NIPBL_geno_F** | GTAGCTACTGTACTGGGTTGTTGTGAG | Primer used for genotyping PCR from upstream of 5' homology arm used for engineering of NIPBL degron cell lines. |
| **NIPBL_geno_R** | CTTAATGAACAGGGCTTCCTGG | Primer used for genotyping PCR from downstream of 3' homology arm used for engineering of NIPBL degron cell lines. |
| **RAD21_geno_F** | GAAATTACGTTTCAGCATGAGATTTGGA | Primer used for genotyping PCR from upstream of 5' homology arm used for engineering of RAD21 degron cell lines. |
| **RAD21_geno_R** | TGACCAGGTGCATAATTTCCCA | Primer used for genotyping PCR from downstream of 3' homology arm used for engineering of RAD21 degron cell lines. |

**Table S2.**

ChIP-seq experiments performed to assess the distribution of cohesin and associated proteins on chromatin. GEO accession for series is GSE277720.

| Accession | Description | Total reads (single + paired) | duplicate reads | Deduplicated reads | % mouse reads |
|---|---|---|---|---|---|
| **GSM8528478** | ChIP-seq: RPE-1 WT input R1 | 164,486,516 | 42,675,896 | 121,810,620 | NA |
| **GSM8528479** | ChIP-seq: RPE-1 WT CTCF ChIP-seq R1 | 167,966,423 | 45,638,438 | 122,327,985 | NA |
| **GSM8528480** | ChIP-seq: NIPBL-A2 DSG/FA input | 80,722,308 | 27,062,751 | 53,659,557 | NA |
| **GSM8528481** | ChIP-seq: RPE-1 NIPBL-A2 (FKBP/mCh+) DSG/FA HA ChIP | 81,536,112 | 38,216,619 | 43,319,493 | NA |
| **GSM8528482** | ChIP-seq: RPE-1 NIPBL-D7 (FKBP/mGL+) input | 57,901,460 | 22,452,326 | 35,449,134 | NA |
| **GSM8528483** | ChIP-seq: RPE-1 NIPBL-D7 (FKBP/mGL+) STAG1 ChIP | 55,840,198 | 38,323,254 | 17,516,944 | NA |
| **GSM8528484** | ChIP-seq: RPE-1 NIPBL-D7 (FKBP/mGL+) STAG2 ChIP | 56,974,279 | 24,864,831 | 32,109,448 | NA |
| **GSM8528485** | ChIP-seq: RPE-1 NIPBL-A2 (FKBP/mCh+) +DMSO 24hr input (~10% NIH3T3 spike-in) | 44,030,205 | 9,308,230 | 34,721,975 | 10.5 |
| **GSM8528486** | ChIP-seq: RPE-1 NIPBL-A2 (FKBP/mCh+) +dTAGv-1 4hr input (~10% NIH3T3 spike-in) | 49,335,574 | 12,478,359 | 36,857,215 | 10.5 |
| **GSM8528487** | ChIP-seq: RPE-1 NIPBL-A2 (FKBP/mCh+) +dTAGv-1 24hr input (~10% NIH3T3 spike-in) | 35,951,480 | 8,619,626 | 27,331,854 | 8.5 |
| **GSM8528488** | ChIP-seq: RPE-1 NIPBL-D7 (FKBP/mGL+) +DMSO 24hr input (~10% NIH3T3 spike-in) | 41,618,461 | 9,130,679 | 32,487,782 | 5.6 |
| **GSM8528489** | ChIP-seq: RPE-1 NIPBL-D7 (FKBP/mGL+) +dTAGv-1 4hr input (~10% NIH3T3 spike-in) | 41,183,151 | 7,809,345 | 33,373,806 | 5.8 |
| **GSM8528490** | ChIP-seq: RPE-1 NIPBL-D7 (FKBP/mGL+) +dTAGv-1 24hr input (~10% NIH3T3 spike-in) | 36,930,826 | 6,618,666 | 30,312,160 | 6.9 |
| **GSM8528491** | ChIP-seq: RPE-1 NIPBL-A2 (FKBP/mCh+) +DMSO 24hr RAD21 (ab992) ChIP (~10% NIH3T3 spike-in) | 30,986,489 | 7,757,883 | 23,228,606 | 14.9 |
| **GSM8528492** | ChIP-seq: RPE-1 NIPBL-A2 (FKBP/mCh+) +dTAGv-1 4hr RAD21 (ab992) ChIP (~10% NIH3T3 spike-in) | 58,726,964 | 14,206,150 | 44,520,814 | 19 |
| **GSM8528493** | ChIP-seq: RPE-1 NIPBL-A2 (FKBP/mCh+) +dTAGv-1 24hr RAD21 (ab992) ChIP (~10% NIH3T3 spike-in) | 52,313,696 | 18,312,579 | 34,001,117 | 30.7 |
| **GSM8528494** | ChIP-seq: RPE-1 NIPBL-D7 (FKBP/mGL+) +DMSO 24hr RAD21 (ab992) ChIP (~10% NIH3T3 spike-in) | 42,178,984 | 12,146,880 | 30,032,104 | 7.6 |
| **GSM8528495** | ChIP-seq: RPE-1 NIPBL-D7 (FKBP/mGL+) +dTAGv-1 4hr RAD21 (ab992) ChIP (~10% NIH3T3 spike-in) | 67,216,428 | 23,401,404 | 43,815,024 | 14.5 |
| **GSM8528496** | ChIP-seq: RPE-1 NIPBL-D7 (FKBP/mGL+) +dTAGv-1 24hr RAD21 (ab992) ChIP (~10% NIH3T3 spike-in) | 51,485,972 | 15,002,231 | 36,483,741 | 19 |
| **GSM8528497** | ChIP-seq: RPE-1 WT input | 45,400,825 | 9,048,788 | 36,352,037 | NA |
| **GSM8528498** | ChIP-seq: RPE-1 WT WAPL (PTG) ChIP-seq | 48,523,707 | 12,299,241 | 36,224,466 | NA |
| **GSM8528499** | ChIP-seq: RPE-1 NIPBL-D7 (FKBP/mGL+) +DMSO 24hr PDS5 ChIP (~10% NIH3T3 spike-in) | 39,470,842 | 11,268,253 | 28,202,589 | 4.2 |

### Table S3.

Hi-C experiments used to determine the consequences of NIPBL and RAD21 depletion on 3D genome organization. GEO accession for series is GSE277719.

| Accession | Description | Total reads (single + paired) | duplicate reads | Deduplicated reads | % cis <1kb | % cis >1kb | % trans |
|---|---|---|---|---|---|---|---|
| **GSM8528442** | Hi-C: RPE-1 NIPBL-A2 (FKBP/mC+) +DMSO 24hr | 662,613,062 | 171,086,946 | 491,526,116 | 11.5 | 79.5 | 9.0 |
| **GSM8528443** | Hi-C: RPE-1 NIPBL-D7 (FKBP/mGL+) +DMSO 24hr | 831,594,994 | 221,268,490 | 610,326,504 | 12.5 | 78.3 | 9.2 |
| **GSM8528444** | Hi-C: RPE-1 NIPBL-A2 (FKBP/mC+) +dTAGv-1 4hr | 652,021,782 | 164,875,200 | 487,146,582 | 18.7 | 71.0 | 10.3 |
| **GSM8528445** | Hi-C: RPE-1 NIPBL-D7 (FKBP/mGL+) +dTAGv-1 4hr | 689,718,315 | 165,692,262 | 524,026,053 | 32.6 | 58.4 | 9.0 |
| **GSM8528446** | Hi-C: RPE-1 NIPBL-A2 (NIPBL-FKBP/mC+) +dTAGv-1 24hr | 758,008,672 | 188,327,045 | 569,681,627 | 12.4 | 75.4 | 12.2 |
| **GSM8528447** | Hi-C: RPE-1 NIPBL-D7 (NIPBL-FKBP/mC+) +dTAGv-1 24hr | 812,028,844 | 213,448,870 | 598,579,974 | 13.4 | 73.6 | 13.0 |
| **GSM8528448** | Hi-C: RPE-1 NIPBL-D7 (FKBP/mGL+) +DMSO, post-nocodazole t=0 R1 | 431,382,852 | 66,229,979 | 365,152,873 | 13.7 | 83.1 | 3.2 |
| **GSM8528449** | Hi-C: RPE-1 NIPBL-D7 (FKBP/mGL+) +dTAG, post-nocodazole t=0 R1 | 563,973,567 | 98,133,749 | 465,839,818 | 14.2 | 81.8 | 4.0 |
| **GSM8528450** | Hi-C: RPE-1 NIPBL-D7 (FKBP/mGL+) +DMSO, post-nocodazole t=30 R1 | 436,269,249 | 73,300,351 | 362,968,898 | 14.0 | 82.7 | 3.3 |
| **GSM8528451** | Hi-C: RPE-1 NIPBL-D7 (FKBP/mGL+) +dTAG, post-nocodazole t=30 R1 | 622,786,308 | 104,009,108 | 518,777,200 | 12.8 | 82.9 | 4.3 |
| **GSM8528452** | Hi-C: RPE-1 NIPBL-D7 (FKBP/mGL+) +DMSO, post-nocodazole t=60 R1 | 319,898,029 | 50,865,584 | 269,032,445 | 15.5 | 78.3 | 6.2 |
| **GSM8528453** | Hi-C: RPE-1 NIPBL-D7 (FKBP/mGL+) +dTAG, post-nocodazole t=60 R1 | 434,118,253 | 76,466,957 | 357,651,296 | 15.8 | 75.9 | 8.3 |
| **GSM8528454** | Hi-C: RPE-1 NIPBL-D7 (FKBP/mGL+) +DMSO, post-nocodazole t=90 R1 | 505,502,814 | 95,828,837 | 409,673,977 | 17.3 | 74.9 | 7.8 |
| **GSM8528455** | Hi-C: RPE-1 NIPBL-D7 (FKBP/mGL+) +dTAG, post-nocodazole t=90 R1 | 376,254,572 | 60,438,772 | 315,815,800 | 16.3 | 73.0 | 10.7 |
| **GSM8528456** | Hi-C: RPE-1 NIPBL-D7 (FKBP/mGL+) +DMSO, post-nocodazole t=120 R1 | 604,740,475 | 123,266,696 | 481,473,779 | 22.4 | 69.2 | 8.4 |
| **GSM8528457** | Hi-C: RPE-1 NIPBL-D7 (FKBP/mGL+) +dTAG, post-nocodazole t=120 R1 | 640,247,666 | 124,257,535 | 515,990,131 | 18.2 | 69.6 | 12.2 |
| **GSM8528458** | Hi-C: RPE-1 NIPBL-D7 (FKBP/mGL+) +DMSO, post-nocodazole t=0 R2 | 479,251,026 | 84,102,938 | 395,148,088 | 14.9 | 82.4 | 2.7 |
| **GSM8528459** | Hi-C: RPE-1 NIPBL-D7 (FKBP/mGL+) +dTAG, post-nocodazole t=0 R2 | 435,371,552 | 73,310,561 | 362,060,991 | 20.7 | 76.6 | 2.7 |
| **GSM8528460** | Hi-C: RPE-1 NIPBL-D7 (FKBP/mGL+) +DMSO, post-nocodazole t=30 R2 | 324,749,887 | 53,126,726 | 271,623,161 | 12.6 | 84.7 | 2.7 |
| **GSM8528461** | Hi-C: RPE-1 NIPBL-D7 (FKBP/mGL+) +dTAG, post-nocodazole t=30 R2 | 498,694,474 | 89,909,475 | 408,784,999 | 15.0 | 81.6 | 3.4 |
| **GSM8528462** | Hi-C: RPE-1 NIPBL-D7 (FKBP/mGL+) +DMSO, post-nocodazole t=60 R2 | 549,522,168 | 98,421,617 | 451,100,551 | 17.5 | 76.9 | 5.6 |
| **GSM8528463** | Hi-C: RPE-1 NIPBL-D7 (FKBP/mGL+) +dTAG, post-nocodazole t=60 R2 | 373,820,884 | 60,426,743 | 313,394,141 | 15.4 | 77.3 | 7.3 |
| **GSM8528464** | Hi-C: RPE-1 NIPBL-D7 (FKBP/mGL+) +DMSO, post-nocodazole t=90 R2 | 493,145,035 | 82,888,718 | 410,256,317 | 18.9 | 74.0 | 7.2 |
| **GSM8528465** | Hi-C: RPE-1 NIPBL-D7 (FKBP/mGL+) +dTAG, post-nocodazole t=90 R2 | 618,920,383 | 137,703,733 | 481,216,650 | 19.1 | 70.8 | 10.1 |
| **GSM8528466** | Hi-C: RPE-1 NIPBL-D7 (FKBP/mGL+) +DMSO, post-nocodazole t=120 R2 | 463,886,608 | 88,993,565 | 374,893,043 | 19.8 | 71.4 | 8.8 |
| **GSM8528467** | Hi-C: RPE-1 NIPBL-D7 (FKBP/mGL+) +dTAG, post-nocodazole t=120 R2 | 451,342,390 | 78,012,103 | 373,330,287 | 19.6 | 68.3 | 12.1 |
| **GSM8528468** | Hi-C: RPE-1 NIPBL-D7 (FKBP/mGL+) +DMSO, post-nocodazole t=8hr R1 | 890,624,410 | 196,707,052 | 693,917,358 | 18.7 | 70.0 | 11.4 |
| **GSM8528469** | Hi-C: RPE-1 NIPBL-D7 (FKBP/mGL+) +dTAG, post-nocodazole t=8hr R1 | 468,620,649 | 104,333,706 | 364,286,943 | 18.5 | 66.2 | 15.4 |
| **GSM8528470** | Hi-C: RPE-1 NIPBL-D7 (FKBP/mGL+) +DMSO, post-nocodazole t=8hr R2 | 932,249,086 | 290,246,333 | 642,002,753 | 19.7 | 68.9 | 11.4 |
| **GSM8528471** | Hi-C: RPE-1 NIPBL-D7 (FKBP/mGL+) +dTAG, post-nocodazole t=8hr R2 | 919,780,691 | 265,868,975 | 653,911,716 | 17.3 | 67.2 | 15.5 |

| | | | | | | | |
|---|---|---|---|---|---|---|---|
| **GSM8528472** | Hi-C: RPE-1 RAD21-B1 (FKBP/mGL+) +DMSO PI sort G1 | 569,168,809 | 254,847,925 | 314,320,884 | 21.3 | 70.1 | 8.5 |
| **GSM8528473** | Hi-C: RPE-1 RAD21-B3 (FKBP/mGL+) +DMSO PI sort G1 | 450,102,748 | 192,505,576 | 257,597,172 | 23.1 | 68.6 | 8.3 |
| **GSM8528474** | Hi-C: RPE-1 RAD21-B1 (FKBP/mGL+) +dTAG 4hr PI sort G1 | 462,156,114 | 177,041,924 | 285,114,190 | 25.8 | 60.7 | 13.5 |
| **GSM8528475** | Hi-C: RPE-1 RAD21-B3 (FKBP/mGL+) +dTAG 4hr PI sort G1 | 339,634,309 | 131,535,258 | 208,099,051 | 25.4 | 60.9 | 13.7 |
| **GSM8528476** | Hi-C: RPE-1 RAD21-B1 (FKBP/mGL+) +dTAG 24hr PI sort G1 | 648,519,561 | 280,396,405 | 368,123,156 | 27.7 | 58.1 | 14.2 |
| **GSM8528477** | Hi-C: RPE-1 RAD21-B3 (FKBP/mGL+) +dTAG 24hr PI sort G1 | 529,887,932 | 218,561,586 | 311,326,346 | 32.4 | 54.4 | 13.2 |

### Table S4.

SLAM-seq experiments performed to determine the consequences of NIPBL depletion on gene expression. GEO accession for series is GSE277898.

| Accession | Description | Total reads (single + paired) | duplicate reads | Deduplicated reads |
|---|---|---|---|---|
| **GSM8533996** | SLAM-seq: RPE-1 NIPBL-D7 (FKBP/mGL+) +dTAG, post-nocodazole t=0, 15min 4sU R1 | 89,808,852 | 31,100,037 | 58,708,815 |
| **GSM8533997** | SLAM-seq: RPE-1 NIPBL-D7 (FKBP/mGL+) +dTAG, post-nocodazole t=30, 15min 4sU R1 | 135,631,067 | 58,030,642 | 77,600,425 |
| **GSM8533998** | SLAM-seq: RPE-1 NIPBL-D7 (FKBP/mGL+) +DMSO, post-nocodazole t=60, 15min 4sU R1 | 154,916,205 | 74,598,277 | 80,317,928 |
| **GSM8533999** | SLAM-seq: RPE-1 NIPBL-D7 (FKBP/mGL+) +dTAG, post-nocodazole t=60, 15min 4sU R1 | 122,835,634 | 47,969,028 | 74,866,606 |
| **GSM8534000** | SLAM-seq: RPE-1 NIPBL-D7 (FKBP/mGL+) +DMSO, post-nocodazole t=90, 15min 4sU R1 | 76,442,853 | 33,870,040 | 42,572,813 |
| **GSM8534001** | SLAM-seq: RPE-1 NIPBL-D7 (FKBP/mGL+) +dTAG, post-nocodazole t=90, 15min 4sU R1 | 71,888,652 | 25,627,325 | 46,261,327 |
| **GSM8534002** | SLAM-seq: RPE-1 NIPBL-D7 (FKBP/mGL+) +DMSO, post-nocodazole t=120, 15min 4sU R1 | 109,771,885 | 54,977,415 | 54,794,470 |
| **GSM8534003** | SLAM-seq: RPE-1 NIPBL-D7 (FKBP/mGL+) +dTAG, post-nocodazole t=120, 15min 4sU R1 | 125,793,019 | 56,936,716 | 68,856,303 |
| **GSM8534004** | SLAM-seq: RPE-1 NIPBL-D7 (FKBP/mGL+) +DMSO, post-nocodazole t=0, 15min 4sU R2 | 143,152,814 | 55,651,310 | 87,501,504 |
| **GSM8534005** | SLAM-seq: RPE-1 NIPBL-D7 (FKBP/mGL+) +dTAG, post-nocodazole t=0, 15min 4sU R2 | 228,465,794 | 105,416,002 | 123,049,792 |
| **GSM8534006** | SLAM-seq: RPE-1 NIPBL-D7 (FKBP/mGL+) +DMSO, post-nocodazole t=30, 15min 4sU R2 | 163,095,418 | 67,181,523 | 95,913,895 |
| **GSM8534007** | SLAM-seq: RPE-1 NIPBL-D7 (FKBP/mGL+) +dTAG, post-nocodazole t=30, 15min 4sU R2 | 144,633,101 | 67,726,905 | 76,906,196 |
| **GSM8534008** | SLAM-seq: RPE-1 NIPBL-D7 (FKBP/mGL+) +DMSO, post-nocodazole t=60, 15min 4sU R2 | 105,851,349 | 37,994,096 | 67,857,253 |
| **GSM8534009** | SLAM-seq: RPE-1 NIPBL-D7 (FKBP/mGL+) +dTAG, post-nocodazole t=60, 15min 4sU R2 | 79,506,171 | 28,078,757 | 51,427,414 |
| **GSM8534010** | SLAM-seq: RPE-1 NIPBL-D7 (FKBP/mGL+) +DMSO, post-nocodazole t=90, 15min 4sU R2 | 130,189,730 | 56,967,405 | 73,222,325 |
| **GSM8534011** | SLAM-seq: RPE-1 NIPBL-D7 (FKBP/mGL+) +dTAG, post-nocodazole t=90, 15min 4sU R2 | 122,259,588 | 61,156,019 | 61,103,569 |
| **GSM8534012** | SLAM-seq: RPE-1 NIPBL-D7 (FKBP/mGL+) +DMSO, post-nocodazole t=120, 15min 4sU R2 | 97,541,957 | 40,306,007 | 57,235,950 |
| **GSM8534013** | SLAM-seq: RPE-1 NIPBL-D7 (FKBP/mGL+) +dTAG, post-nocodazole t=120, 15min 4sU R2 | 132,299,001 | 59,951,283 | 72,347,718 |
| **GSM8534014** | SLAM-seq: RPE-1 NIPBL-D7 (FKBP/mGL+) +DMSO, post-nocodazole t=0, 15min 4sU R3 | 155,329,099 | 122,512,162 | 32,816,937 |
| **GSM8534015** | SLAM-seq: RPE-1 NIPBL-D7 (FKBP/mGL+) +dTAG, post-nocodazole t=0, 15min 4sU R3 | 114,136,127 | 46,235,672 | 67,900,455 |
| **GSM8534016** | SLAM-seq: RPE-1 NIPBL-D7 (FKBP/mGL+) +DMSO, post-nocodazole t=30, 15min 4sU R3 | 73,743,269 | 25,948,606 | 47,794,663 |
| **GSM8534017** | SLAM-seq: RPE-1 NIPBL-D7 (FKBP/mGL+) +dTAG, post-nocodazole t=30, 15min 4sU R3 | 86,369,489 | 31,776,533 | 54,592,956 |
| **GSM8534018** | SLAM-seq: RPE-1 NIPBL-D7 (FKBP/mGL+) +DMSO, post-nocodazole t=60, 15min 4sU R3 | 137,378,592 | 91,773,304 | 45,605,288 |
| **GSM8534019** | SLAM-seq: RPE-1 NIPBL-D7 (FKBP/mGL+) +dTAG, post-nocodazole t=60, 15min 4sU R3 | 126,309,785 | 75,668,640 | 50,641,145 |
| **GSM8534020** | SLAM-seq: RPE-1 NIPBL-D7 (FKBP/mGL+) +DMSO, post-nocodazole t=90, 15min 4sU R3 | 89,254,530 | 48,637,547 | 40,616,983 |
| **GSM8534021** | SLAM-seq: RPE-1 NIPBL-D7 (FKBP/mGL+) +dTAG, post-nocodazole t=90, 15min 4sU R3 | 144,240,966 | 68,628,340 | 75,612,626 |
| **GSM8534022** | SLAM-seq: RPE-1 NIPBL-D7 (FKBP/mGL+) +DMSO, post-nocodazole t=120, 15min 4sU R3 | 117,472,615 | 99,851,782 | 17,620,833 |
| **GSM8534023** | SLAM-seq: RPE-1 NIPBL-D7 (FKBP/mGL+) +DMSO, post-nocodazole t=0, 15min 4sU R4 | 79,260,897 | 26,121,809 | 53,139,088 |
| **GSM8534024** | SLAM-seq: RPE-1 NIPBL-D7 (FKBP/mGL+) +dTAG, post-nocodazole t=0, 15min 4sU R4 | 91,776,753 | 31,558,030 | 60,218,723 |
| **GSM8534025** | SLAM-seq: RPE-1 NIPBL-D7 (FKBP/mGL+) +DMSO, post-nocodazole t=30, 15min 4sU R4 | 132,117,668 | 61,809,242 | 70,308,426 |

| | | | | |
|---|---|---|---|---|
| **GSM8534026** | SLAM-seq: RPE-1 NIPBL-D7 (FKBP/mGL+) +DMSO, post-nocodazole t=60, 15min 4sU R4 | 136,384,383 | 61,060,850 | 75,323,533 |
| **GSM8534027** | SLAM-seq: RPE-1 NIPBL-D7 (FKBP/mGL+) +dTAG, post-nocodazole t=60, 15min 4sU R4 | 106,609,323 | 42,072,595 | 64,536,728 |
| **GSM8534028** | SLAM-seq: RPE-1 NIPBL-D7 (FKBP/mGL+) +DMSO, post-nocodazole t=90, 15min 4sU R4 | 114,139,674 | 77,651,982 | 36,487,692 |
| **GSM8534029** | SLAM-seq: RPE-1 NIPBL-D7 (FKBP/mGL+) +dTAG, post-nocodazole t=90, 15min 4sU R4 | 138,994,563 | 59,352,490 | 79,642,073 |
| **GSM8534030** | SLAM-seq: RPE-1 NIPBL-D7 (FKBP/mGL+) +DMSO, post-nocodazole t=120, 15min 4sU R4 | 82,740,706 | 33,996,744 | 48,743,962 |
| **GSM8534031** | SLAM-seq: RPE-1 NIPBL-D7 (FKBP/mGL+) +dTAG, post-nocodazole t=120, 15min 4sU R4 | 108,365,435 | 37,096,440 | 71,268,995 |
| **GSM8534032** | SLAM-seq: RPE-1 NIPBL-D7 (FKBP/mGL+) +DMSO, post-nocodazole t=120, 120min 4sU R1 | 89,189,834 | 34,388,174 | 54,801,660 |
| **GSM8534033** | SLAM-seq: RPE-1 NIPBL-D7 (FKBP/mGL+) +dTAG, post-nocodazole t=120, 120min 4sU R1 | 115,788,213 | 46,519,265 | 69,268,948 |
| **GSM8534034** | SLAM-seq: RPE-1 NIPBL-D7 (FKBP/mGL+) +DMSO, post-nocodazole t=120, 120min 4sU R2 | 89,285,697 | 32,724,608 | 56,561,089 |
| **GSM8534035** | SLAM-seq: RPE-1 NIPBL-D7 (FKBP/mGL+) +dTAG, post-nocodazole t=120, 120min 4sU R2 | 80,122,837 | 28,247,469 | 51,875,368 |
| **GSM8534036** | SLAM-seq: RPE-1 NIPBL-D7 (FKBP/mGL+) +DMSO, post-nocodazole t=120, 120min 4sU R3 | 89,659,259 | 34,527,626 | 55,131,633 |
| **GSM8534037** | SLAM-seq: RPE-1 NIPBL-D7 (FKBP/mGL+) +dTAG, post-nocodazole t=120, 120min 4sU R3 | 42,713,975 | 12,773,851 | 29,940,124 |
| **GSM8534038** | SLAM-seq: RPE-1 NIPBL-D7 (FKBP/mGL+), post-nocodazole 4hr release +DMSO t=30min, 15min 4sU R1 | 107,125,233 | 66,745,249 | 40,379,984 |
| **GSM8534039** | SLAM-seq: RPE-1 NIPBL-D7 (FKBP/mGL+), post-nocodazole 4hr release +dTAG t=30min, 15min 4sU R1 | 197,611,894 | 153,371,706 | 44,240,188 |
| **GSM8534040** | SLAM-seq: RPE-1 NIPBL-D7 (FKBP/mGL+), post-nocodazole 4hr release +DMSO t=60min, 15min 4sU R1 | 126,029,108 | 95,582,200 | 30,446,908 |
| **GSM8534041** | SLAM-seq: RPE-1 NIPBL-D7 (FKBP/mGL+), post-nocodazole 4hr release +dTAG t=60min, 15min 4sU R1 | 131,612,127 | 105,731,013 | 25,881,114 |
| **GSM8534043** | SLAM-seq: RPE-1 NIPBL-D7 (FKBP/mGL+), post-nocodazole 4hr release +DMSO t=90min, 15min 4sU R1 | 81,658,288 | 41,427,664 | 40,230,624 |
| **GSM8534045** | SLAM-seq: RPE-1 NIPBL-D7 (FKBP/mGL+), post-nocodazole 4hr release +dTAG t=90min, 15min 4sU R1 | 140,335,158 | 105,488,873 | 34,846,285 |
| **GSM8534047** | SLAM-seq: RPE-1 NIPBL-D7 (FKBP/mGL+), post-nocodazole 4hr release +DMSO t=120min, 15min 4sU R1 | 144,960,286 | 98,058,135 | 46,902,151 |
| **GSM8534049** | SLAM-seq: RPE-1 NIPBL-D7 (FKBP/mGL+), post-nocodazole 4hr release +dTAG t=120min, 15min 4sU R1 | 161,591,549 | 136,152,094 | 25,439,455 |
| **GSM8534052** | SLAM-seq: RPE-1 NIPBL-D7 (FKBP/mGL+), post-nocodazole 4hr release +DMSO t=30min, 15min 4sU R2 | 129,339,227 | 66,561,120 | 62,778,107 |
| **GSM8534053** | SLAM-seq: RPE-1 NIPBL-D7 (FKBP/mGL+), post-nocodazole 4hr release +dTAG t=30min, 15min 4sU R2 | 161,464,419 | 93,673,025 | 67,791,394 |
| **GSM8534054** | SLAM-seq: RPE-1 NIPBL-D7 (FKBP/mGL+), post-nocodazole 4hr release +DMSO t=60min, 15min 4sU R2 | 236,423,716 | 183,830,325 | 52,593,391 |
| **GSM8534055** | SLAM-seq: RPE-1 NIPBL-D7 (FKBP/mGL+), post-nocodazole 4hr release +dTAG t=60min, 15min 4sU R2 | 124,660,897 | 65,149,251 | 59,511,646 |
| **GSM8534056** | SLAM-seq: RPE-1 NIPBL-D7 (FKBP/mGL+), post-nocodazole 4hr release +DMSO t=90min, 15min 4sU R2 | 134,229,823 | 72,093,314 | 62,136,509 |
| **GSM8534057** | SLAM-seq: RPE-1 NIPBL-D7 (FKBP/mGL+), post-nocodazole 4hr release +dTAG t=90min, 15min 4sU R2 | 140,991,667 | 66,403,567 | 74,588,100 |
| **GSM8534058** | SLAM-seq: RPE-1 NIPBL-D7 (FKBP/mGL+), post-nocodazole 4hr release +DMSO t=120min, 15min 4sU R2 | 169,234,109 | 102,229,013 | 67,005,096 |
| **GSM8534059** | SLAM-seq: RPE-1 NIPBL-D7 (FKBP/mGL+), post-nocodazole 4hr release +dTAG t=120min, 15min 4sU R2 | 133,023,237 | 74,776,840 | 58,246,397 |
| **GSM8534060** | SLAM-seq: RPE-1 NIPBL-D7 (FKBP/mGL+), post-nocodazole 4hr release +DMSO t=30min, 15min 4sU R3 | 132,134,913 | 89,463,377 | 42,671,536 |
| **GSM8534061** | SLAM-seq: RPE-1 NIPBL-D7 (FKBP/mGL+), post-nocodazole 4hr release +dTAG t=30min, 15min 4sU R3 | 125,487,793 | 112,496,597 | 12,991,196 |
| **GSM8534062** | SLAM-seq: RPE-1 NIPBL-D7 (FKBP/mGL+), post-nocodazole 4hr release +DMSO t=60min, 15min 4sU R3 | 122,766,171 | 93,302,330 | 29,463,841 |
| **GSM8534063** | SLAM-seq: RPE-1 NIPBL-D7 (FKBP/mGL+), post-nocodazole 4hr release +dTAG t=60min, 15min 4sU R3 | 172,536,225 | 111,926,842 | 60,609,383 |
| **GSM8534064** | SLAM-seq: RPE-1 NIPBL-D7 (FKBP/mGL+), post-nocodazole 4hr release +DMSO t=90min, 15min 4sU R3 | 165,753,554 | 131,101,645 | 34,651,909 |
| **GSM8534065** | SLAM-seq: RPE-1 NIPBL-D7 (FKBP/mGL+), post-nocodazole 4hr release +dTAG t=90min, 15min 4sU R3 | 136,516,277 | 63,625,650 | 72,890,627 |

| | | | | |
|---|---|---|---|---|
| **GSM8534066** | SLAM-seq: RPE-1 NIPBL-D7 (FKBP/mGL+), post-nocodazole 4hr release +DMSO t=120min, 15min 4sU R3 | 140,356,335 | 74,658,255 | 65,698,080 |
| **GSM8534067** | SLAM-seq: RPE-1 NIPBL-D7 (FKBP/mGL+), post-nocodazole 4hr release +dTAG t=120min, 15min 4sU R3 | 123,404,375 | 50,903,774 | 72,500,601 |
| **GSM8534068** | SLAM-seq: RPE-1 NIPBL-D7 (FKBP/mGL+), post-nocodazole 4hr release +DMSO t=120min, 120min 4sU R1 | 143,852,474 | 94,867,343 | 48,985,131 |
| **GSM8534069** | SLAM-seq: RPE-1 NIPBL-D7 (FKBP/mGL+), post-nocodazole 4hr release +dTAG t=120min, 120min 4sU R1 | 131,047,862 | 76,412,353 | 54,635,509 |
| **GSM8534070** | SLAM-seq: RPE-1 NIPBL-D7 (FKBP/mGL+), post-nocodazole 4hr release +DMSO t=120min, 120min 4sU R2 | 203,047,898 | 101,966,984 | 101,080,914 |
| **GSM8534071** | SLAM-seq: RPE-1 NIPBL-D7 (FKBP/mGL+), post-nocodazole 4hr release +dTAG t=120min, 120min 4sU R2 | 47,860,502 | 14,691,644 | 33,168,858 |
| **GSM8534072** | SLAM-seq: RPE-1 NIPBL-D7 (FKBP/mGL+), post-nocodazole 4hr release +DMSO t=120min, 120min 4sU R3 | 104,460,943 | 49,924,463 | 54,536,480 |
| **GSM8534073** | SLAM-seq: RPE-1 NIPBL-D7 (FKBP/mGL+), post-nocodazole 4hr release +dTAG t=120min, 120min 4sU R3 | 79,460,354 | 32,400,953 | 47,059,401 |
| **GSM8534074** | SLAM-seq: RPE-1 NIPBL-D7 (FKBP/mGL+) +DMSO, post-nocodazole t=4hr, 15min 4sU R1 | 254,759,857 | 155,153,369 | 99,606,488 |
| **GSM8534075** | SLAM-seq: RPE-1 NIPBL-D7 (FKBP/mGL+) +dTAG, post-nocodazole t=4hr, 15min 4sU R1 | 120,516,558 | 55,460,560 | 65,055,998 |
| **GSM8534076** | SLAM-seq: RPE-1 NIPBL-D7 (FKBP/mGL+) +DMSO, post-nocodazole t=6hr, 15min 4sU R1 | 134,576,735 | 61,636,996 | 72,939,739 |
| **GSM8534077** | SLAM-seq: RPE-1 NIPBL-D7 (FKBP/mGL+) +dTAG, post-nocodazole t=6hr, 15min 4sU R1 | 113,366,360 | 54,754,700 | 58,611,660 |
| **GSM8534078** | SLAM-seq: RPE-1 NIPBL-D7 (FKBP/mGL+) +DMSO, post-nocodazole t=8hr, 15min 4sU R1 | 159,354,601 | 83,787,725 | 75,566,876 |
| **GSM8534079** | SLAM-seq: RPE-1 NIPBL-D7 (FKBP/mGL+) +dTAG, post-nocodazole t=8hr, 15min 4sU R1 | 93,243,827 | 37,953,269 | 55,290,558 |
| **GSM8534080** | SLAM-seq: RPE-1 NIPBL-D7 (FKBP/mGL+) +DMSO, post-nocodazole t=4hr, 15min 4sU R2 | 184,406,133 | 103,209,359 | 81,196,774 |
| **GSM8534081** | SLAM-seq: RPE-1 NIPBL-D7 (FKBP/mGL+) +dTAG, post-nocodazole t=4hr, 15min 4sU R2 | 92,072,304 | 38,783,774 | 53,288,530 |
| **GSM8534082** | SLAM-seq: RPE-1 NIPBL-D7 (FKBP/mGL+) +DMSO, post-nocodazole t=6hr, 15min 4sU R2 | 136,558,539 | 58,036,570 | 78,521,969 |
| **GSM8534083** | SLAM-seq: RPE-1 NIPBL-D7 (FKBP/mGL+) +dTAG, post-nocodazole t=6hr, 15min 4sU R2 | 103,449,342 | 39,250,904 | 64,198,438 |
| **GSM8534084** | SLAM-seq: RPE-1 NIPBL-D7 (FKBP/mGL+) +DMSO, post-nocodazole t=8hr, 15min 4sU R2 | 179,039,399 | 91,397,571 | 87,641,828 |
| **GSM8534085** | SLAM-seq: RPE-1 NIPBL-D7 (FKBP/mGL+) +dTAG, post-nocodazole t=8hr, 15min 4sU R2 | 137,104,667 | 66,710,506 | 70,394,161 |

**Table S5.**

ChIP-seq datasets accessed from the Sequence Read Archive (SRA).

| Description | SRA | Run | Control | Application |
|---|---|---|---|---|
| **H3K36me3 ChIP-seq** | SRP359812 | SRR23280340 | SRR18024435 | ChromHMM |
| **H3K27me3 ChIP-seq** | SRP359812 | SRR23280342 | SRR18024435 | ChromHMM |
| **H3K9ac ChIP-seq** | SRP337334 | SRR15871778 | SRR15871781 | ChromHMM |
| **H3k4me3 ChIP-seq** | SRP298142 | SRR13259862 | SRR13259866 | ChromHMM |
| **H3k4me1 ChIP-seq** | SRP298142 | SRR13259864 | SRR13259866 | ChromHMM |
| **H3K9me3 ChIP-seq** | SRP337334 | SRR15871779 | SRR15871781 | ChromHMM |
| **H3K27ac ChIP-seq** | SRP233381 | SRR10540139 | SRR10540157 | ChromHMM, ROSE |
| **H3K27ac ChIP-seq** | SRP233381 | SRR10540140 | SRR10540160 | ROSE |
| **SMC3ac ChIP-seq** | SRP359812 | SRR18024438 | SRR18024437 | Persistent chromatin loops |
